# A mathematical and computational model of the calcium dynamics in *Caenorhabditis elegans* ASH sensory neuro

**DOI:** 10.1101/201962

**Authors:** Ehsan Mirzakhalili, Bogdan Epureanu, Eleni Gourgou

**Author notes:** E.G. current address: Mechanical Engineering Dept., University of Michigan, Ann Arbor, MI. corresponding author: Eleni Gourgou.

## Abstract

We propose a mathematical and computational model that captures the stimulus-generated Ca^2+^ transients in the *C. elegans* ASH sensory neuron. The model is built based on biophysical events and molecular cascades known to unfold as part of neurons’ Ca^2+^ homeostasis mechanism, as well as on Ca^2+^ signaling events. The state of ion channels is described by their probability of being activated or inactivated, and the remaining molecular states are based on biochemically defined kinetic equations with phenomenological adjustments. We estimate the parameters of the model using experimental data of hyperosmotic stimulus-evoked Ca^2+^ transients detected with a FRET sensor in young and aged worms, unstressed and exposed to oxidative stress. We use a hybrid optimization method composed of a multi-objective genetic algorithm and nonlinear least-squares to estimate the model parameters. We first obtain the model parameters for young unstressed worms. Next, we use these values of the parameters as a starting point to identify the model parameters for stressed and aged worms. We show that the model, in combination with experimental data, corroborates literature results. In addition, we demonstrate that our model can be used to predict ASH response to complex combinations of stimulation pulses. The proposed model includes for the first time the ASH Ca^2+^ dynamics observed during both "on" and "off" responses. This mathematical and computational effort is the first to propose a dynamic model of the Ca^2+^ transients’ mechanism in *C. elegans* neurons, based on biochemical pathways of the cell’s Ca^2+^ homeostasis machinery.

**Significance Statement:** *C. elegans* is widely used as a model system for monitoring neuronal Ca^2+^ transients. The ASH neuron is the subject of several such studies, primarily due to its key importance as a polymodal nociceptor. However, despite its pivotal role in *C. elegans* biology, and the special characteristics of its stimulus-evoked Ca^2+^ transients (e.g., the "off" response), no mathematical or computational model has been developed to include special features of ASH Ca^2+^ dynamics, i.e. the "off" response. The model includes for the first time the ASH Ca^2+^ dynamics observed during both "on" and "off" responses, and is the first to propose a dynamical model of the *C. elegans* Ca^2+^ transients’ mechanism based on biochemical pathways of the cell’s Ca^2+^ homeostasis machinery.

**Abbreviations:**

ER: endoplasmic reticulum
PMCA: plasma membrane Ca^2+^ ATPase
SERCA: sarco-endoplasmic reticulum Ca^2+^ -transport ATPase
TRPV: transient receptor potential-vallinoid
VGCC: voltage gated Ca^2+^ channels
IP_3_: 3-phopsho inositol
IPR: IP_3_ receptors
ROS: reactive oxygen species
GA: genetic algorithm
ES: extracellular space

## Introduction

The use of Ca^2+^ transients to indirectly assess a neuron’s activation is a well-established approach (1–3) despite its limitations, and despite the extra care needed to draw conclusions about the neuron’s concurrent activation (4–7). A the same time, *C. elegans* has been proved ideal for applying imaging techniques to monitor Ca^2+^ transients generated upon stimulation of a variety of neurons (2, 8–13), in freely moving (9, 14) as well as in immobilized worms, by using traditional approaches (3, 4, 15, 16) or advanced methods (2, 17–21). The ASH polymodal neuron is the subject of numerous such studies (17, 18, 21, 22), because of its key importance as a nociceptor for the worms’ survival and also because it is the starting player for a plethora of downstream neuronal events. The use of microfluidic chips for the immobilization of worms and the delivery of stimuli has revealed not only the first peak in the Ca^2+^ transients of the ASH neuron occurring upon stimulation (the "on" response), but also a second peak occurring upon withdrawal of the stimulus (the "off" response) (18, 23, 24).

The bimodal Ca^2+^ currents that occur in the ASH neuron upon its stimulation are the object of several studies, which explore the connection between Ca^2+^ transients, neuronal behavior (6, 23–25) and synaptic output of the ASH neuron to downstream neurons (10, 18, 26). In particular, the connection between Ca^2+^ transients in the ASH neuron and *C. elegans* behavior has been the object of several studies that suggest an interesting correlation not only between the "on" response and specific behaviors (27, 29), but also between the "off" response and avoidance behavior (23). This leads to the conclusion that all features of the Ca^2+^ dynamics in the ASH neuron participate in fine-tuning the worm’s rich behavioral repertoire.

In parallel, Ca^2+^ transients (3) have been studied in ASH neurons in the context of different biological conditions, including aging (17, 22), oxidative stress (11, 21), food availability (29) and oxygen concentration (30). Extended efforts have been made to decipher the molecular players involved (3, 4, 23, 25, 31). At the same time, mathematical modeling of Ca^2+^ dynamics has been performed in a variety of organisms and cells, but the literature is sparse and far from complete for mathematical modeling of Ca^2+^ response in *C. elegans* neurons (27, 32, 33). Work conducted by Kato and colleagues (24) has focused on the temporal responses of both ASH and AWC to flickering stimuli, developing a phenomenological model that explains selected features of Ca^2+^ dynamics using ordinary differential equations. However, that work does not include the ASH "off" response, and the model does not account for the dynamics of the molecular players involved.

We propose a mathematical and computational model that is based on biochemical pathways and for the first time encompasses the Ca^2+^ dynamics observed during both "on" and "off" responses in the ASH neuron in *C. elegans.* The approach integrates biophysical models that describe several aspects of the overall Ca^2+^ signaling mechanism and it merges them into an inclusive model forged by novel phenomenological adjustments. Thus, the model succeeds to capture the Ca^2+^ dynamics in ASH neurons with excellent fidelity. Moreover, the model can be used to suggest potential changes in molecular components that can explain modifications in Ca^2+^ dynamics due to aging and oxidative stress. Lastly, we demonstrate how the proposed model can be used to predict Ca^2+^ transients in ASH neuron when delivering arrays of complex stimuli.

## Methods

The model is based on intracellular molecular events responsible for the generation of Ca^2+^ transients. Numerous molecules and pathways are involved in this process (34, 35). We focused the model on molecular players which are believed to dominate the dynamics (4, 24, 36-39), taking also into account that some of the remaining molecular players are not yet well understood in *C. elegans* neurons (40). We modeled the dynamics of the secondary players using a coarse grain approach, similar to reduced order modeling. To this end, we introduced selected equivalent players (viewed as states in the model) whose dynamics captures the overall combined effects of most of the secondary players. The pathways and molecules we include in the proposed model are depicted in Figure 1. A detailed description of their contribution in the generation of Ca^2+^ transients and their connection with the rest of the players in the model is given in the Discussion.

**Figure 1:**
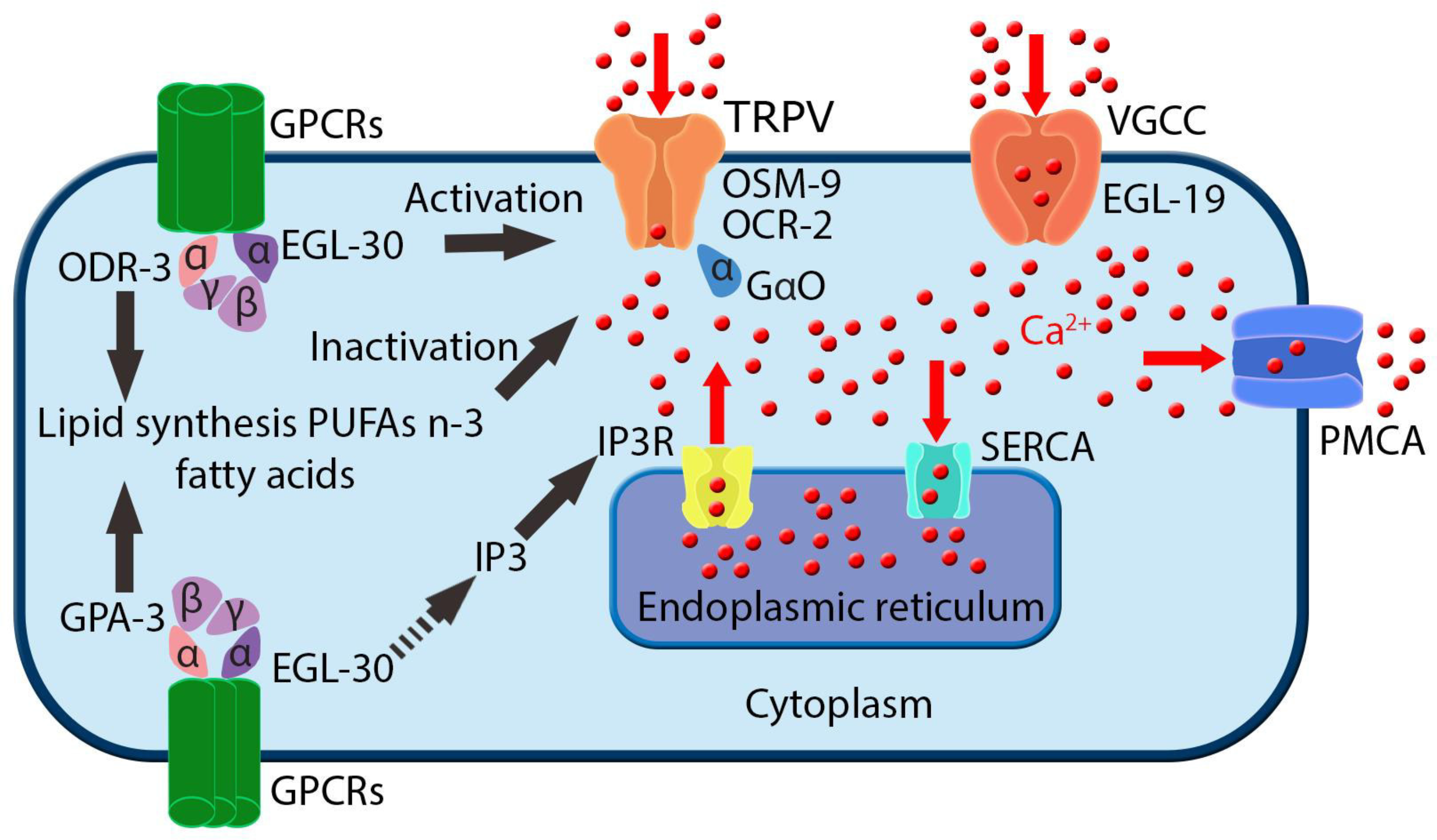
Molecular components of the Ca^2+^ homeostatic machinery that are included in the proposed mathematical model. GPCRs: G-protein coupled receptors, as the ones that are activated in ASH neuron by hyperosmotic stimuli; ODR-3, EGL-30, GPA-3: G proteins coupled with the receptors, participating in signal transduction to downstream ion channels; OSM-9, OCR-2: molecular elements of the TRPV channels, the main cation channels through which Ca^2+^ flows into the neuron upon its stimulation; GaO: G-protein coupled with TRPVs; EGL-19: molecular component of the VGCCs, the L-type voltage gated Ca^2+^ channels, activated by the changed membrane potential due to ion influx upon neuronal activation; PMCA: plasma membrane Ca^2+^ ATPase, the main pump responsible for transporting Ca^2+^ into the extracellular space; SERCA: sarco-endoplasmic reticulum Ca^2+^ ATPase, which transports Ca^2+^ into the intracellular stores; IP_3_: 3-phopsho-inositol, secondary messenger participating in Ca^2+^ signaling events; IP_3_R: IP_3_ receptors, glycoprotein complex acting as a Ca2+ channel activated by IP_3_, abundant on the endoplasmic reticulum (ER) membranes. GPCRs, ODR-3, EGL-30, GPA-3, OSM-9, OCR-2: not modeled individually; model parameters that account for these molecular components are P_0_, P_1_ and P_2_, see Methods. Lipid synthesis, PUFAs, fatty acids, TRPV activation, TRPV inactivation events: modeled as O (activated) and I (inactivated) probabilities, see Methods. Included in the model and not depicted here: J_LEAK_ and J_LEAK,ER_, which represent the constant influx of Ca^2+^ into the cytoplasm from extracellular space and ER, respectively, through other mechanisms, see Methods.

### Mathematical Model of Ca^2+^ Dynamics

Assuming a cell with well-mixed free Ca^2+^, the concentration of free Ca^2+^ in the cytoplasm and endoplasmic reticulum (ER) can be written as the following system of ordinary differential equations (41):

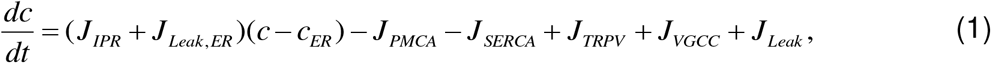

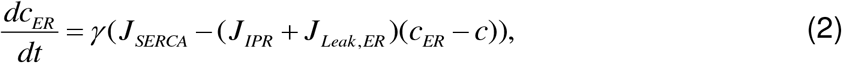

Where *c* and *c*_*ER*_ are the concentrations of free Ca^2+^ in the cytoplasm and in the ER. *J*_*PMCA*_ represents the Ca^2+^ efflux through plasma membrane Ca^2+^ ATPase (PMCA) pumps to the extracellular space (ES), and *J*_*SERCA*_ is the Ca^2+^ flux from the cytoplasm into ER through ER ATPase pumps. *J*_*TRPV*_ and *J*_*VGCC*_ denote the influxes of Ca^2+^ into cytoplasm from ES through transient receptor potential-vallinoid (TRPV) channels and through voltage gated Ca^2+^ channels (VGCC). *J*_*Leak*_ and *J*_*Leak*,*ER*_ represent constant influxes of Ca^2+^ into the cytoplasm from ES and ER through other mechanisms. *γ* denotes the ratio of the cytoplasmic volume to the ER volume, which can also account for fast linear Ca^2+^ buffers in the ER (42).

The PMCA pumps are modeled following (41) as:

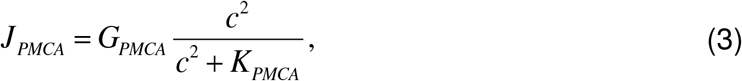

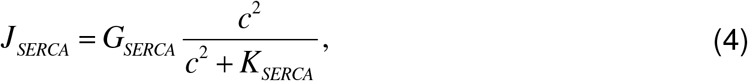

where *G*_*PMCA*_ and *G*_*SERCA*_ are the maximum fluxes through PMCA and SERCA pumps. *K*_*PMCA*_ and *K*_*SERCA*_ model the affinity of PMCA and SERCA for Ca^2+^.

The Ca^2+^ influx through TRPV channels is modeled as:

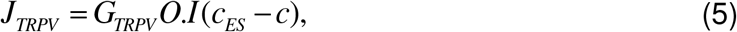

where *G*_*TRPV*_ is the maximum influx of Ca^2+^ through TRPV channels, and *c*_*ES*_ denotes the Ca^2+^ concentration in the ES, which is assumed to be constant. *O* and *I* are the probabilities of TRPV channels to be activated and inactivated, and are governed by:

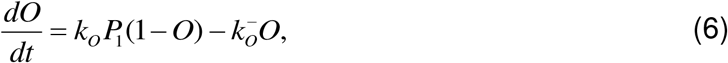

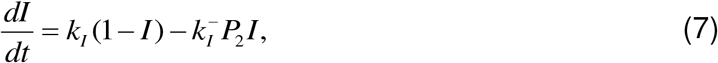

where *k*_*O*_, *k*_*I*_ are forward and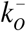, 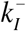 are backward rate constants for the *O* and *I* states. *P*_1_ and *P*_2_ represent two molecular players that control activation and inactivation of TRPV channels with the following kinetics:

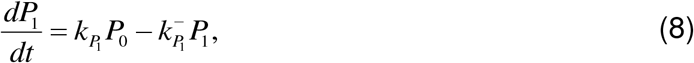

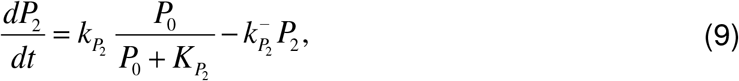

where 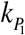, 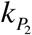 are forward and 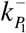, 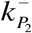 are backward rate constants for *P*_1_ and *P*_2_, and 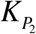 is the affinity of *P*_2_ for an equivalent player *P*_0_ that coarsely represents the dynamic effects of the cascade of molecular players that are activated by the external stimulus.

We model the dynamics of *P*_0_ as:

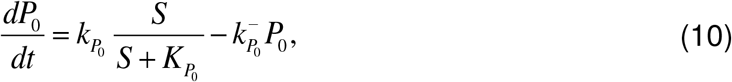

where *S* represents the strength of the stimulus, 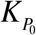 is the affinity of *P*_0_ for *S*, and 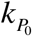, 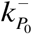 are forward and backward rates for *P*_0_.

Player *P*_0_ coarsely accounts for the pathway that is triggered upon delivery of the stimulus, including the G-proteins which are coupled with the receptors (43, 44) (ODR-3, EGL-30, Figure 1). Player *P*_0_ coarsely represents the pathway downstream of EGL-30, which leads to activation of TRPV channels (43). Player *P*_2_ represents the cascade of events in which fatty acids participate, modulating TRPV channels (35, 45) (Figure 1).

The Ca^2+^ influx through IP3 receptors (IP3Rs) is modeled as:

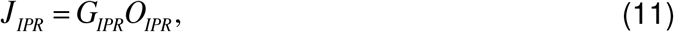

where *G*_*IPR*_ is the maximum Ca^2+^ flux through IP_3_ receptors, and *O*_*IPR*_ is the probability of IP_3_ receptors (IPRs) to be open, which is modeled using a reduced form (46, 47) in the De Young-Keizer model (48), namely:

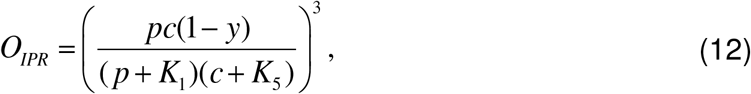

where *p* is the concentration of IP_3_, and *y* is the faction of inhibited IPRs. The faction of inhibited IPRs is in turn governed by:

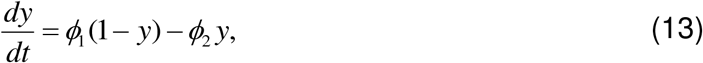

where

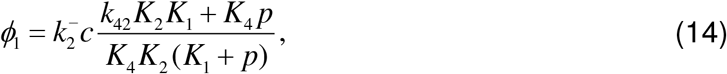

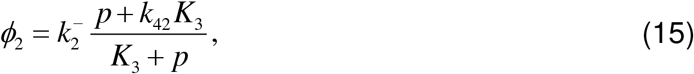

with 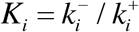 being the equilibrium constants for binding/unbinding of IP_3_ or Ca^2+^ to IPRs, with their original values, as used by De Young and Keizer (48).

To model variations in *p* upon the delivery of the stimulus, we use players *P*_1_ and *P*_2_ together with *c* to create the following phenomenological model:

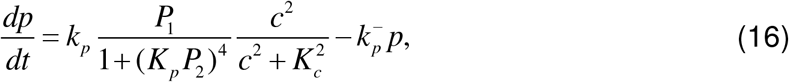

where *k*_*p*_ and 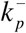 are constants that represent forward and backward rates, while *K*_*p*_ and *K*_*c*_ are the affinities of *p* for *P*_2_ and *c*. The relation between *P*_1_ and *P*_2_ in Eq. (16) follows an incoherent feed-forward network motif (49), which is also combined with a Hill equation (41) for Ca^2+^. This part of the model plays an important role in the "off" response.

The Ca^2+^ influx through L-type voltage gated calcium channels (VGCCs) are modeled using the Goldman-Hodgkin-Katz (50) equation:

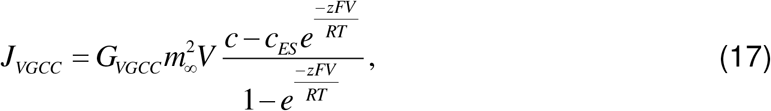

where *G*_*VGCC*_ is the maximum Ca^2+^ flux through VGCC, *z* is the valence of the respective ions ( *z* = 2 for Ca^2+^), ***F*** is the Faraday constant, ***R*** is the gas constant, *T* is the temperature, *V* is the membrane voltage, and `m` is the activation variable given by:

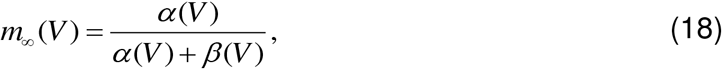

where *α* and *β* are rates that vary with the membrane voltage for the L-type current as follows (51):

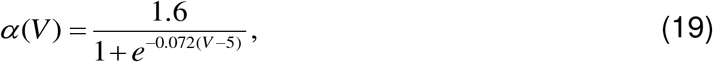

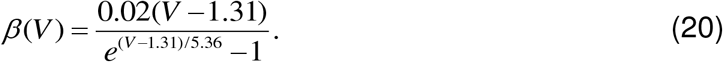

Finally, we model the dynamics of membrane potential *V*. Unfortunately, the available experimental data regarding changes in the membrane potential of the *C. elegans* ASH neuron is limited. Moreover, the ion currents, which are responsible for the voltage response of *C. elegans* neurons, are not completely understood. However, the available data suggests that many *C. elegans* neurons show a graded voltage response rather than the (better characterized and more commonly encountered) action potential response (52–54). Therefore, we use a phenomenological approach to model a graded voltage response based on the only ionic current, namely the Ca^2+^ current, as follows:

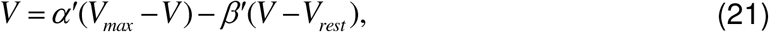

where *V*_*max*_ is the maximum membrane potential that can be reached during the graded response of the ASH neuron, and *V*_*rest*_ = −70mV is the resting membrane potential. *α*′ and *β*′ are forward and backward rates for Eq. (21), and are given by:

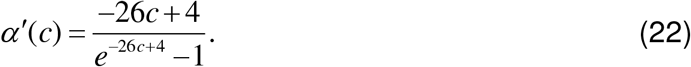

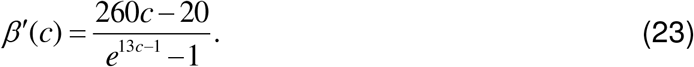

Note that *α′* and *β′* are similar to *α* and *β* in Eqs. (19) and (20). However, they have different values, and are functions of *c* (concentration of free Ca^2+^ in the cytoplasm) instead of *V*.

### Ca^2+^ Concentration to FRET Signal Conversion

The Ca^2+^ concentration obtained using the mathematical model can be related to signals measured in experiments (17, 21). Experimental signals obtained using TN-XL FRET (fluorescence resonance energy transfer) measurements (55) are the result of Ca^2+^ interaction with an indicator genetically encoded in the *C. elegans* ASH neuron. The relation between the Ca^2+^ concentration and the measured % FRET change signal ***R*** follows an empirical form (56) that can be expressed as:

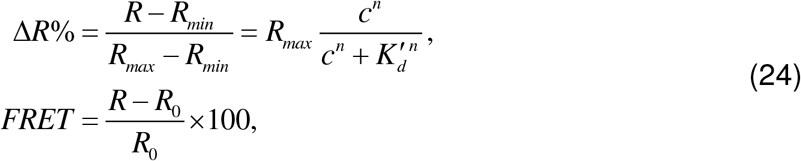

where the Hill coefficient is *n* = 1.7, and the apparent affinity of ***R*** for Ca^2+^ is 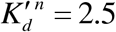, as provided in (55).

Equation (24) can be used to convert measured % FRET changes into the corresponding Ca^2+^ concentration. Using Eq. (22) requires the values of *R*_*min*_, *R*_*max*_ and *R*_*0*_. To this end, we used the data in (57, 58). We set the baseline Ca^2+^ concentration to 100nM and the maximum Ca^2+^ concentration to 500nM when Δ*R*% = 10 (for young unstressed worms), and then calculated *R*_*min*_ and *R*_*max*_. Next, we determined *R*_0_ using Eq. (24) based on the initial value of the Ca^2+^ concentration in the mathematical model, which is set to 100nM for all cases.

### Parameter estimation

There are 23 parameters (Table S1) that need to be determined in order to use the mathematical model proposed. Ultimately, the mathematical model has to capture the Ca^2+^ dynamics in four experimental data sets that are used in this study, namely measured Ca^2+^ transients for young unstressed, young stressed, aged unstressed and aged stressed worms. We chose the experimental data set for young unstressed worms as reference to determine all 23 parameters. Next, we modified the values of as few of these parameters as necessary to fit the data for the other three cases, one at a time. This approach allows us to suggest different possible pathways through which the parameters governing Ca^2+^ transients may change, to capture alterations in stimulus-evoked Ca^2+^ dynamics in aged worms or in worms previously exposed to oxidative stress.

We used a hybrid optimization approach consisting of a genetic algorithm (GA) as a global minimizer, and trust-region-reflective nonlinear least squares (TRNLS) as a local optimizer, to determine the parameters of the mathematical model. The hybrid optimization algorithm starts with the GA, and the fittest individual found from the GA is passed to the TRNLS. The minimization problem is also subjected to constraints that ensure all parameter values are physical (namely that they are positive). The MATLAB optimization toolbox is used to perform these calculations. The maximum-likelihood of the experimental data is the objective function used in the TRNLS algorithm, namely:

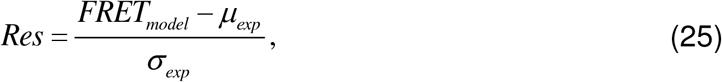

where *Res* is the residual at each measurement instant, *FRET*_*model*_ is the data obtained from the model, *μ*_*exp*_ and *σ*_*exp*_ are the average and the standard deviation of experimental data at that same time instant. The objective function to be minimized in the GA is the sum of all residuals given by Eq. (25) over all time instants measured.

We determined first the parameters for young unstressed worms. The solution of the GA depends on the initial population. Thus, we first applied the GA as a multi-objective optimizer to all four sets of experimental data. We then used the resulting population as an initial population for the GA applied only to the data from the young unstressed worms. Among the optimum solutions suggested by the multi-objective GA, we chose the solution for which the sum of the residuals for all experimental cases was the minimum, even though there were other solutions in which the residual for individual cases was smaller than the chosen solution. This approach allowed us to select as initial population for the GA a vicinity in the parameter space that is near an optimum solution (minimum residual) for all four cases.

There are more than one parameter sets that can result in the measured Ca^2+^ dynamics for young unstressed worms. However, we sought a single set of parameters for young unstressed worms for which the sensitivity of the solution for the estimated parameters was small. To that end, we used the numerical Jacobian matrix that is obtained from TRNLS to approximate the Hessian matrix. Then, we used the Hessian matrix to construct the covariance matrix. We chose the solution that had the smallest diagonal elements of the covariance matrix. That corresponds to choosing the solution with the smallest variance of the parameters, indicating that small changes in the parameters (for that solution) do not lead to vast changes in the Ca^2+^ dynamics.

For the other three experimental cases, we start with the parameters found for young unstressed worms to create the initial population for the GA in the hybrid optimization. Next, we limit the algorithm to change only 13 selected parameters of the total of 23. These 13 parameters (listed with bold in Supplementary Table 1) include strength and rates related to key players of the Ca^2+^ signaling mechanism – TRPV channels being activated/deactivated, IP_3_, IPRs, PMCAs, SERCAs – and they have been selected based on discussions in the existing literature (59–62). Then, we use the hybrid optimization algorithm to estimate the values of the 13 selected parameters, while the other parameters are kept constant. A similar procedure is performed separately for all possible combinations for the selected parameters. Next, the results from all the combinations are pooled together for each of the three experimental cases. Each combination of selected parameters is a potential pathway that can show effects of aging or oxidative stress on young unstressed worms. However, not all of the combinations are plausible. The first criterion used to select valid combinations of parameter sets, is the goodness of fit that the mathematical model provides, i.e. the residuals must be small. To make the comparison consistent among different cases, we sort the residual for all different combinations and only keep the combinations for which the residuals are smaller than 99% of all solutions. Next, we use a second criterion on the remaining combinations in which parameter changes that are detectable are selected. To that end, we compute changes in parameters as compared to young unstressed worms. If the absolute value of the changes for a parameter combination is larger than the sensitivity found using the covariance matrix, then we consider such combination plausible.

#### Ca^2+^ transients experimental data analysis

All experimental results presented in Figures 2 and 3 were acquired as described in Gourgou and Chronis, 2016 (21). Briefly, using the TN-XL FRET sensor, the stimulus-evoked Ca^2+^ transients generated when the ASH neuron of stressed (exposed to oxidative stress) and unstressed *C. elegans* of various ages was stimulated by hyperosmotic solution of 1M glycerol (17, 21) were recorded (21). Experimental data used in this study come from four populations of adult hermaphrodite *C. elegans:* i) young (L4+1/Day 1 of adult life) unstressed animals (control animals; reference case), ii) young (L4+1/Day 1 of adult life) oxidative-stressed animals, iii) aged (L4+5/Day 5 of adult life) unstressed animals, and iii) aged (L4+5/Day 5 of adult life) oxidative-stressed animals. It is noted that Day 5 worms can be considered of middle age (63), but for brevity they are referred to as the aged worms.

**Figure 2:**
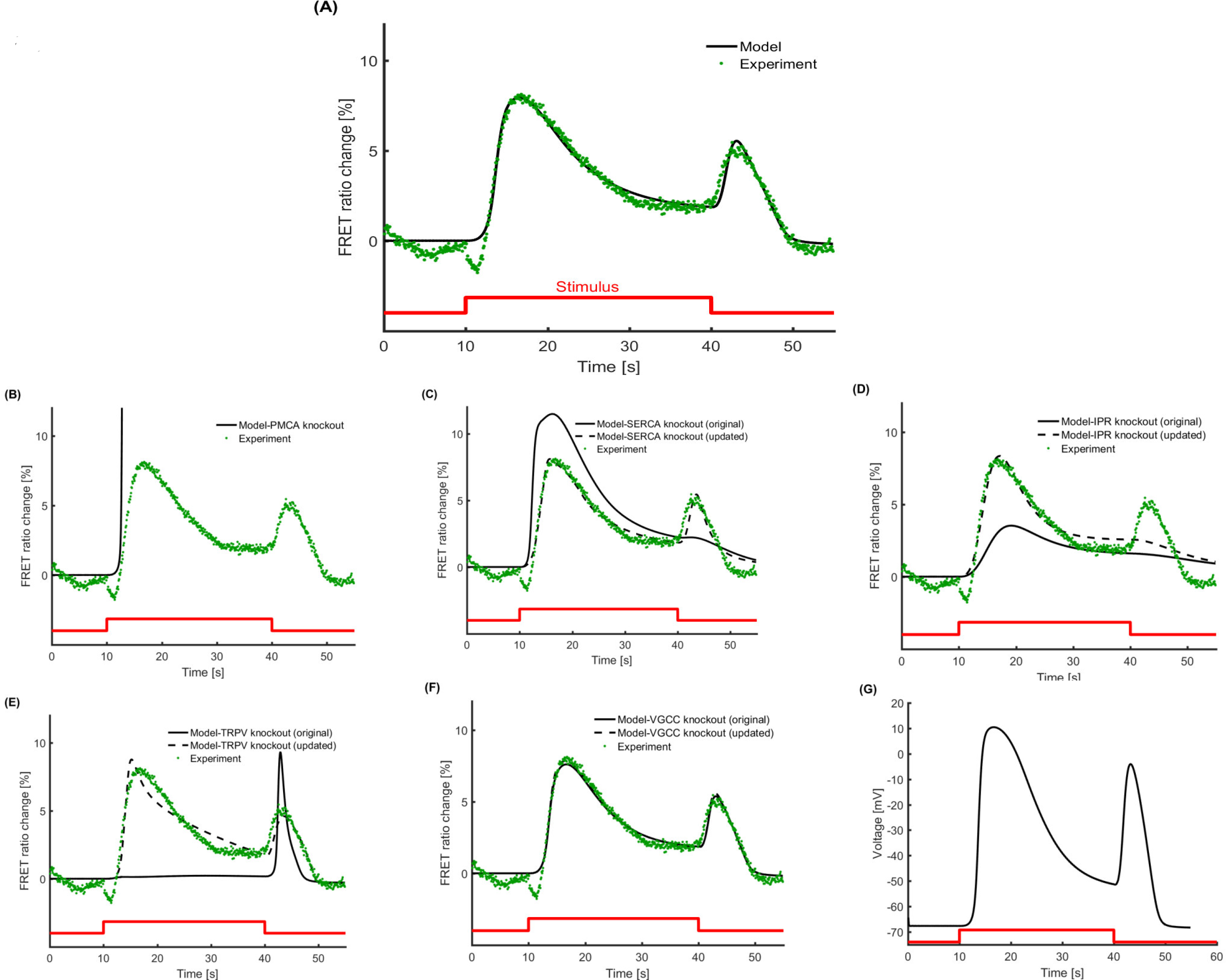
The mathematical model can capture the Ca^2+^ dynamics observed in young (Dayl) unstressed worms, when ASH sensory neuron is stimulated by hyperosmotic solution (glycerol 1M). (A) The mathematical model matches the Ca^2+^ transients, as recorded experimentally, including all key features (time and magnitude of peaks, rising and decay slopes) for the "on" response (upon delivery of the stimulus) and the "off" response (upon withdrawal of the stimulus). (B-F) Different components of the model are knocked out (*in silico k*nock-out) to investigate their impact on the model-generated results. For the knockout results with original parameters (solid line), all the parameters are kept the same as in (A), except for the knockout component that is removed from the mathematical model. For the knockout results with updated parameters (dashed line), the hybrid optimization algorithm is run again trying to find an updated set of parameters which can explain the experimental Ca^2+^ dynamics, after the knockout has been removed from the mathematical model. (B) When PMCA is removed, Ca^2+^ is not pumped out of the cell and the mathematical model fails. Changes in the parameters in any of the model components cannot compensate for PMCA knockout in the updated model. (C) When SERCA is removed, the original model fails to show the "off" response and features of the "on" response are also affected. Updating the parameters restores most of Ca^2+^ transients’ features, except for the decaying slopes. (D) Removal of IPR changes all the features of "on" response and the "off" response completely vanishes. Updating the parameters restores the "on" response partly but fails to return the "off" response. (E) TRPV knockout for the original model does not show any "on" response, while "off" response is amplified. When the parameters are updated, both "on" and "off" responses are partially restored. (F) VGCC *in silico* knock-out does not affect significantly the results of the model, original or updated. (G) The phenomenological model used to describe the voltage response shows a graded response when the stimulus is applied. A weaker graded voltage response is also observed when the stimulus is removed.

**Figure 3:**
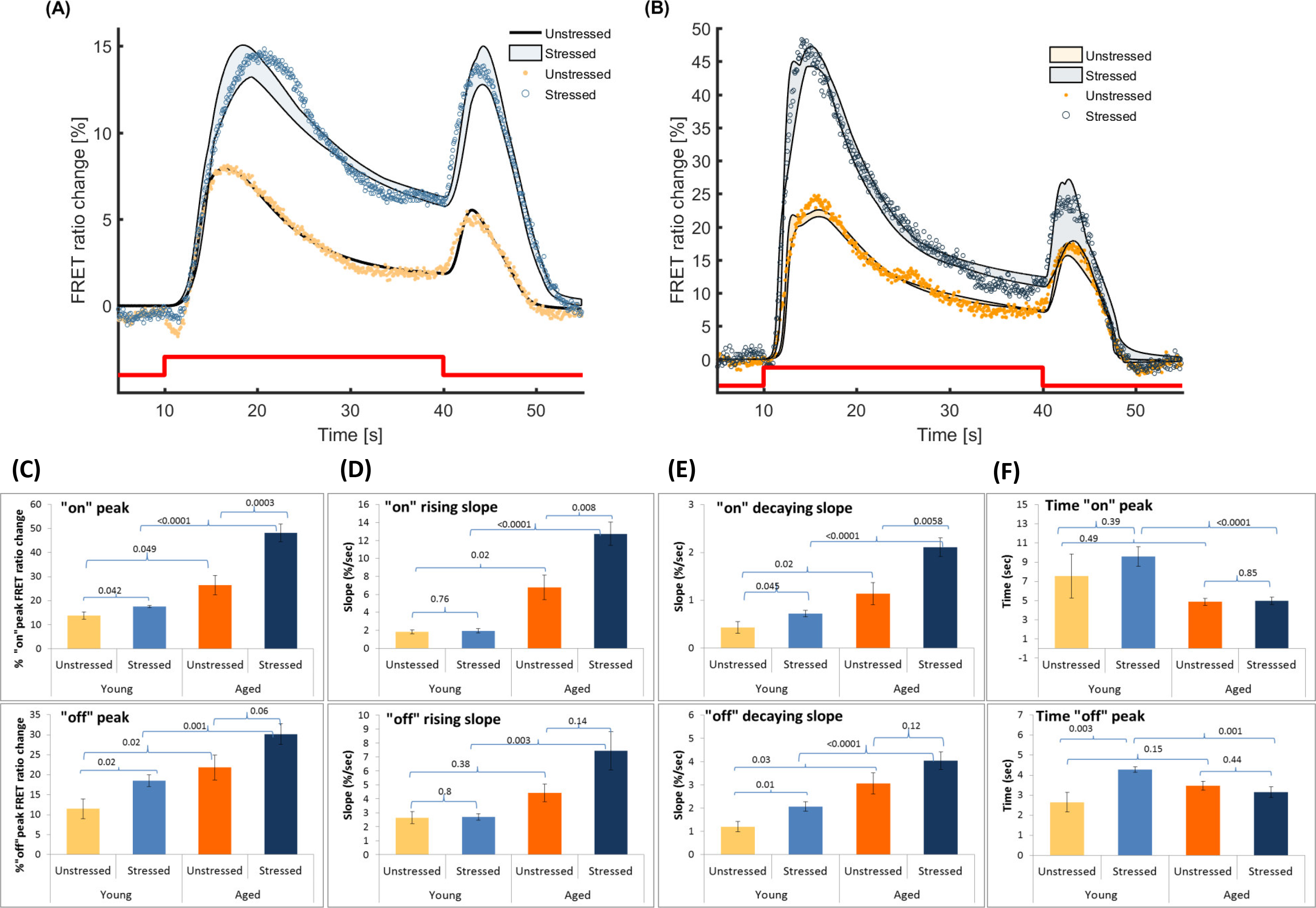
The mathematical model can capture the stimulus-induced changes in the Ca^2+^ dynamics in the case of aged (Day 5) or previously exposed to oxidative stress (stressed) animals. (A) The parameter set for young (Day 1) unstressed worms is used as a reference point to detect changes in the parameters that can explain the stimulus-evoked Ca^2+^ dynamics in treated worms of the same age. The model results shown correspond to all plausible solutions. (B) Similar to (A), the parameter set for young (Day 1) unstressed worms is used to detect changes in the parameters that can explain the stimulus-evoked Ca^2+^ dynamics in aged (Day 5) unstressed and stressed worms. The modeling results correspond to all plausible solutions. Red line represents the stimulus pulse delivered (duration: 30sec). Experimental data modified from Gourgou and Chronis, 2016. (C-F) Key features of the Ca^2+^ transients, as recorded experimentally and presented in (A) and (B); (C) the peak of the "on" (top) and "off" (bottom) response, (D) the rising slope of the "on" (top) and the "off" (bottom) response, (E) the decaying slope of the "on" (top) and the "off" (bottom) response, (F) the time needed to reach the peak of "on" (top) and "off" (bottom) response. Error bars indicate standard error of mean, p-values of Student t-test shown in chart area. Representative results of 17-35 individual experiments. (C) top and (D) top modified from Gourgou and Chronis, 2016.

The peak and the slope of the rising phase of the "on" response in the ratio traces were calculated at the onset of stimulus (Figure 3C, top; see also Figure 2B of Gourgou and Chronis, 2016(21)). The peak of the rising phase is indicative of the total amount of Ca^2+^ entering the cell and consequently accounts for the amplitude of the cell’s response to the applied stimulus, whereas the slope corresponds to the time rate of the Ca^2+^ influx.

Here, we calculate also the decaying slope of the "on" response (Figure 3E, top), which corresponds to the time rate of the Ca^2+^ efflux off the cytoplasm that takes place after the neuron’s initial response to the applied stimulus and leads to a stabilization (plateau) of the cytoplasmic Ca^2+^ after 30 sec of the response onset and until the stimulus is withdrawn. The maximum peak of the "off" response (Figure 3C, bottom) is indicative of the total amount of Ca^2+^ entering the cytoplasm upon withdrawal of the stimulus. The rising and descending slopes of the "off" response (Figures 3D and 3E, bottom panels) account for the time rate of Ca^2+^ influx to the cytoplasm and the time rate of Ca^2+^ efflux off the cytoplasm, both upon withdrawal of the stimulus.

The decaying slope of the "on" response can be defined as the difference between the maximum value of the FRET ratio change on_*max*_ observed upon delivery of the stimulus and the average stabilized FRET signal A_*plateau*_ at the plateau between 30 and 40 sec over the time needed for this namely (on_*max*_ – *A*_*plateau*_) / (*T*_*minplateau*_ – *T*_*max*_), where *T*_*max*_ is the time when on_*max*_ is observed, and *T*_*minplateau*_ is the time when the plateau starts.

The maximum peak *off*_*max*_ of the "off" response is calculated as the % peak FRET ratio change upon withdrawal of the stimulus. The rising slope of the "off" response is the difference between the average stabilized FRET signal A_*plateau*_ at the plateau between 30 and 40 sec and the maximum peak *off*_*max*_ observed upon withdrawal of the stimulus, over the time needed to reach this maximum, counting from the stimulus offset. The rising slope can be expressed as (*off*_*max*_ – *A*_*plateau*_)/(*T*_*max*_ –40).

The decaying slope of the "off" response is the maximum value of the % FRET ratio change *off*_*max*_ observed upon withdrawal of the stimulus over the time needed to reach the minimum value observed after the stimulus offset, namely *off*_*max*_ /(*T*_*offmax*_ – *T*_*min*_).

**Statistical analyses**: All comparisons in Figure 3 are made using two tailed, unpaired Student’s *t*-test. Statistically significant differences were considered the ones with *p*-value < 0.05. Exact *p*-values are provided for each comparison on the respective plot. Statistical analyses were performed in Excel (Microsoft, WA, USA) and Minitab (Minitab Inc., PA, USA).

## Results

### Accuracy and effects of knocking out components of the model

As a first step, we explore whether our model can capture the dynamics of the Ca^2+^ transients which occur when the ASH neuron of young unstressed worms is stimulated by a hyperosmotic solution (Figure 2A). Our results show that the model matches well the special and critical features of the Ca^2+^ transients, including magnitude and time of peaks, rising and decaying slopes for both "on" and "off" responses.

To check if the model is minimal, we removed different components of the model to investigate their contribution to the predicted Ca^2+^ dynamics (Figure 2B-2F). For each of these *in silico* knockouts, two cases were considered: i) the original case, in which after the knockout component is removed the values for all parameters remain the same as in the model that generates the output shown in Figure 2A, and ii) the updated case, in which after the knockout component is removed, we apply anew the hybrid optimization algorithm to estimate again the model parameters.

We find that the model completely diverges without the PMCA pump (Figure 2B), since all the Ca^2+^ that flows in upon delivery of the stimulus, remains in the neuron. Moreover, since the PMCA pump is the only component in the model that actively removes excessive Ca^2+^ outside the neuron, changing the values of any other parameter cannot compensate for the PMCA knockout; therefore, no updated solution can be reached when the PMCA is knocked out.

When the SERCA pump is removed (Figure 2C), Ca^2+^ that enters the cytoplasm is not pumped into the ER. The original model with a knocked out SERCA accounts for more Ca^2+^ in the cytoplasm compared to the results shown in Figure 2A, while at the same time the "off" response is absent. In the updated model, where the parameters of other components are adjusted, the overall results are improved, however the final return of cytoplasmic Ca^2+^ to its initial levels is still not completely restored, and the rising slope of the "off" response is captured with a slight time lag.

Removing IPRs from the model (Figure 2D) completely alters the features of the Ca^2+^ transients. The magnitude of the "on" response decreases dramatically, and the "off" response vanishes. The updated parameters can recover features of the "on" response, but they fail to yield the "off" response.

The "on" response completely disappears when TRPV channels are knocked out (Figure 2E). Interestingly, the "off" response is present and very strong in the absence of TRPV channels. In the updated case, the adjusted parameters yield the "on" response, even without TRPV channels. However, the updated model without the TRPV channels accounts poorly for the magnitude of the "on" response and, most importantly, it fails to capture the dynamics of its decaying slope.

Elimination of VGCCs from the model (Figure 2F) does not impact the model predictions substantially. This is reflected equally in the outputs of the original and the updated version of the model, as shown in Figure 2G.

### The model captures Ca^2+^ dynamics in aged and stressed worms

The values of the parameters in one parameter set found to capture the dynamics of Ca^2+^ for young unstressed worms (Figure 2A) are used as reference from which we can obtain different plausible parameter values that explain Ca^2+^ transients for young stressed (Figure 3A), aged unstressed and aged stressed worms (Figure 3B).

We use the parameters for young unstressed worms to initiate the search for plausible solutions in the other three cases so that we can explore the impact of oxidative stress and age on the model parameters. The experimental data for young stressed worms lie within the bounds of the results predicted by the mathematical model, except for the time right after the "on" response, as well as some of the time on the plateau (Figure 3A), where the model seems not to follow exactly the stabilization in Ca^2+^ concentration. In the case of aged worms, experimental data falls within the bounds of the results predicted by the mathematical model with plausible parameter sets for both unstressed and stressed worms (Figure 3B). Most of the variations generated by the model for aged worms occur around the "off" response, for both unstressed and stressed aged animals.

In parallel, we quantify key features of experimentally recorded Ca^2+^ transients (Figures 3C-3F, top and bottom panels) to identify how they change due to oxidative stress and aging, and eventually correlate them with the model parameters. Results show that the peak of the "on" response increases in stressed conditions for young animals (Figure 3C, bottom). The rising slope of the "on" response increases due to aging, and is affected by oxidative stress only in aged worms (Figure 3D, top). The rising slope of the "off" response increases when both factors, aging and stress, apply (Figure 3D, bottom). Similarly to the peak of the "on" response, the decaying slope of the "on" response increases in stressed and in aged animals (Figure 3E, top). The decaying slope of the "off" response is affected by aging, whereas oxidative stress affects the decaying slope of the "off" response only in young worms (Figure 3E, bottom). Lastly, the "on" response (Figure 3F, top) reaches its peak faster in aged stressed animals, while the "off" response peak occurs earlier in young stressed worms, and later in aged stressed worms (Figure 3F, bottom).

### Parameters that contribute to the modified Ca^2+^ dynamics: Model sensitivity

Each of the plausible parameter sets obtained from the multitude of possible initial populations in the hybrid optimization contains combinations of parameters that are different compared to the reference case (young unstressed worms). The number of times each one of the parameters is present in one of the plausible parameter sets for young stressed (total of 15 plausible sets), aged unstressed (total of 3 plausible sets) and stressed (total of 6 plausible sets) worms is shown in Figures 4A, 4C, and 4E. The absence of a bar for a parameter (for example, VGCC is not present in Figure 4A) does not indicate that the specific parameter was not changed due to the condition studied (i.e., age and/or oxidative stress). Rather, it means that none of the different combinations that constitute plausible solutions for the specific case (e.g., regarding Figure 4A, stress in young worms), contains the absent parameter. As shown in the dot plots in Figures 4B, 4D, and 4F, the values of each parameter in all the plausible sets in which it is included may vary substantially (e.g., *k*_*O*_, Figure 4F) or not (e.g., *k*_*p*_, Figure 4B). Notably, in the case of aged stressed worms (Figure 4F) the parameters included in the plausible solutions that are changed compared to young unstressed worms, are increased hundreds of times (e.g., 4000% for 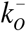, 6000% or even 14000% for *k*_*o*_ Figure 4F). In the case of young stressed and aged unstressed worms, the altered parameters increase only by up to ~350% (Figures 4B, 4D), and sometimes they even decrease, compared to young unstressed animals (e.g. *G*_*SERCA*_ and 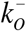 Figure 4B, *K*_*SERCA*_ and *k*_*I*_ in Figure 4D).

**Figure 4:**
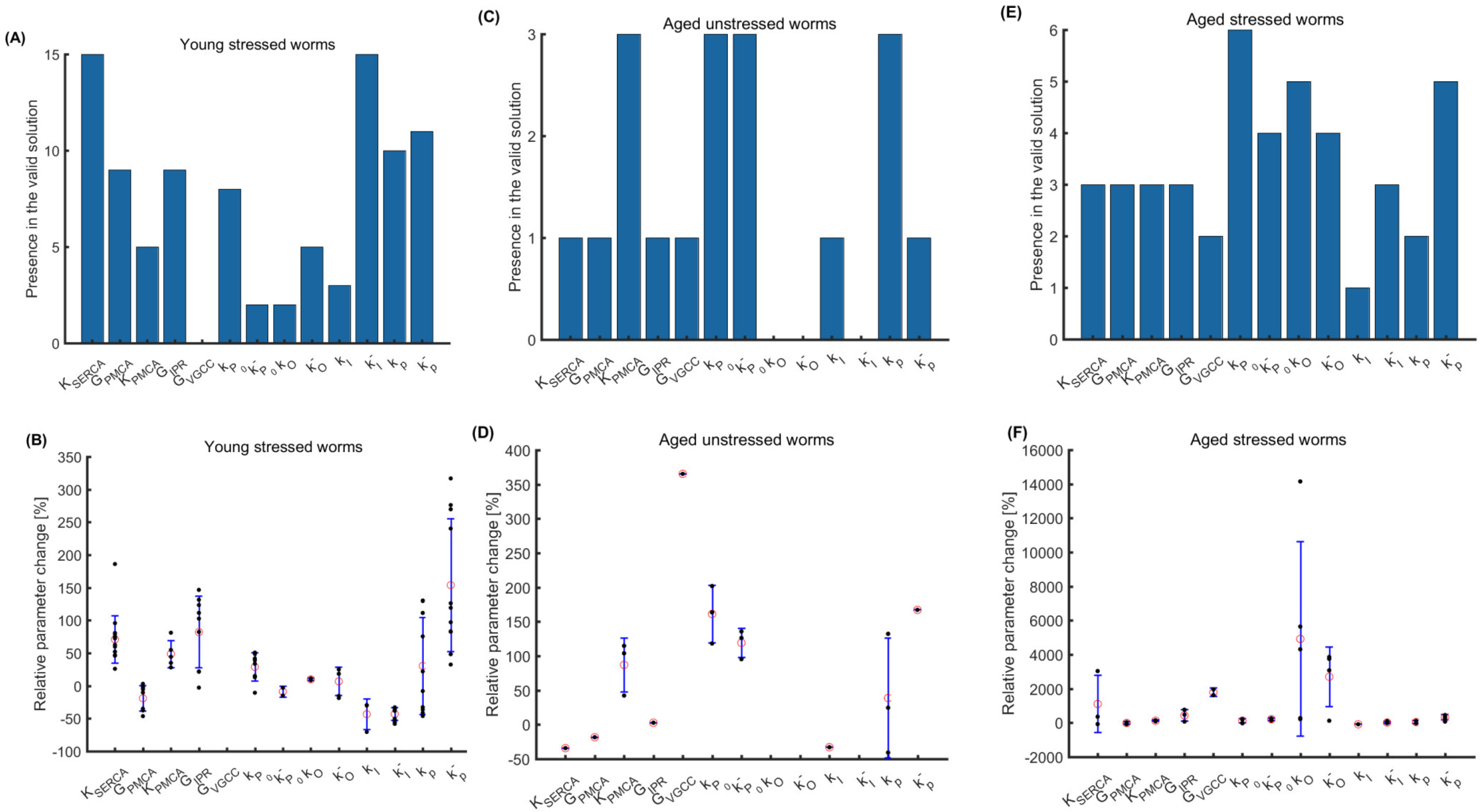
The effects of aging and oxidative stress treatment on stimulus-evoked Ca^2+^ transients can be explained by changing values of the parameter set for young unstressed worms (reference case). The frequency by which each of the selected parameters appears in all plausible combinations of solutions is shown in (A) for young stressed worms (15 plausible combinations), in (C) for aged unstressed worms (3 plausible combinations), and in (E) for aged stressed worms (6 plausible combinations). The dot plots in (B), (D), and (F) show the relative changes in the parameters compared to the respective parameters for young untreated worms. Each dot corresponds to a plausible solution; red circles indicate the mean; error bars represent standard deviation.

The overall sensitivity of the model parameters can be visualized by a sensitivity plot (Figure 5). For each of the plausible parameter sets, the parameter values are randomly perturbed by ±25% in this sensitivity analysis, and the "on" and "off" responses peaks are plotted. Results suggest that the model is able to predict the different behaviors observed in experiments because the model results and the experimental results approximately overlap. Moreover, model-generated data points are clustered near the regions where the experimental data points are dense, and are sparse where the experimental results are scarce. This illustrates that the model can generate more results in the regions where most experimental results are recorded.

**Figure 5:**
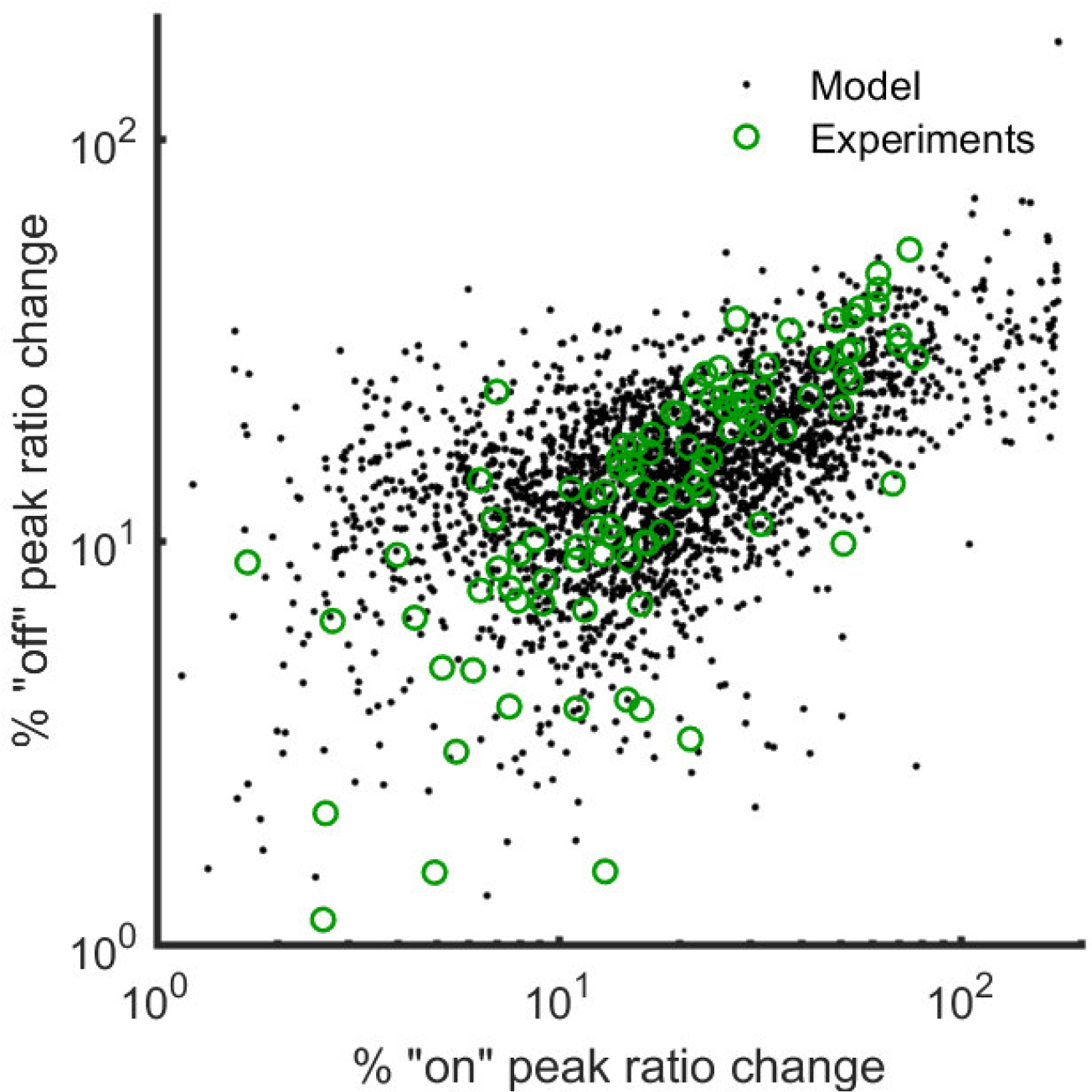
Sensitivity of transients on parameter models. Open green circles show all of the experimental results including young unstressed, young stressed, aged unstressed, and aged stressed worms. For each plausible parameter set 100 samples are created, in which the parameter values are randomly perturbed by ±25%. For each sample, the magnitude maximum "on" and "off" responses are recorded and plotted as filled black circles. The distribution of randomly perturbed model shows that the mathematical model can capture variations that are observed in experimental results. Moreover, the modeling results are observed to be dense/sparse where the experimental results are dense/sparse.

### Using the model to predict Ca^2+^ dynamics in the case of complex stimuli

Next, we used the model to explore how the ASH neuron would respond when activated by complex time-varying stimuli that would be challenging to implement in an experimental setup, yet it is possible for the worms to encounter in nature. Figure 6 demonstrates selected examples of stimulus-generated Ca^2+^ transients for such *in silico* experiments. In all panels, in addition to the %FRET ratio change, the Ca^2+^ concentration in the ER is also presented in order to show the long-lasting effects of the stimulus on the system. Compared to the Ca^2+^ concentration in the cytoplasm (represented by %FRET ratio change), the Ca^2+^ concentration in the ER has a slower dynamic which affects the system response especially during multiple stimuli. At rest, the Ca^2+^ concentration in the ER is equal to its equilibrium value. When the stimulus is delivered, a small increase in Ca^2+^ concentration in the ER is observed due to influx of calcium through TRPV and VGCC, which is then followed by rapid large decrease due to release of Ca^2+^ from the ER through IPR. Then, the Ca^2+^ concentration in the ER remains relatively constant during the stimulus since the influx and efflux of Ca^2+^ balance each other. Finally, another rapid large decrease in the Ca^2+^ concentration in the ER is observed during the "off" response. When the stimulus is withdrawn completely, the Ca^2+^ concentration in the ER starts to go back to its equilibrium. However, if a stimulus is delivered before the Ca^2+^ concentration in the ER reaches its equilibrium (this is the case for all the panels in Figures 6), we observe the effects of previous stimuli on the dynamics of the Ca^2+^ concentration in the cytoplasm. For instance, in all cases with sequential stimuli (Figures 6A-6D), the first "on" response is the strongest. When stimuli of equal strength and different durations are applied (10, 30 and 50 sec) (Figure 6A), the magnitude for all "off" responses is almost the same. The shortest stimuli (fourth and sixth pulse, 10sec) result in just one peak. The longest stimulus (fifth pulse, 50 sec) results in a longer plateau, but similar to the other cases. When we apply stimuli with the same duration but different magnitudes (2α, α, α/2, 4α) (Figure 6B), the response to the first pulse, which has twice the magnitude of the first pulse in Figure 6A, leads to stronger "on" and "off" responses. However, a stimulus with the same magnitude that is delivered later (fourth pulse), following three pulses, it leads to weaker "on" and "off" responses.

**Figure 6:**
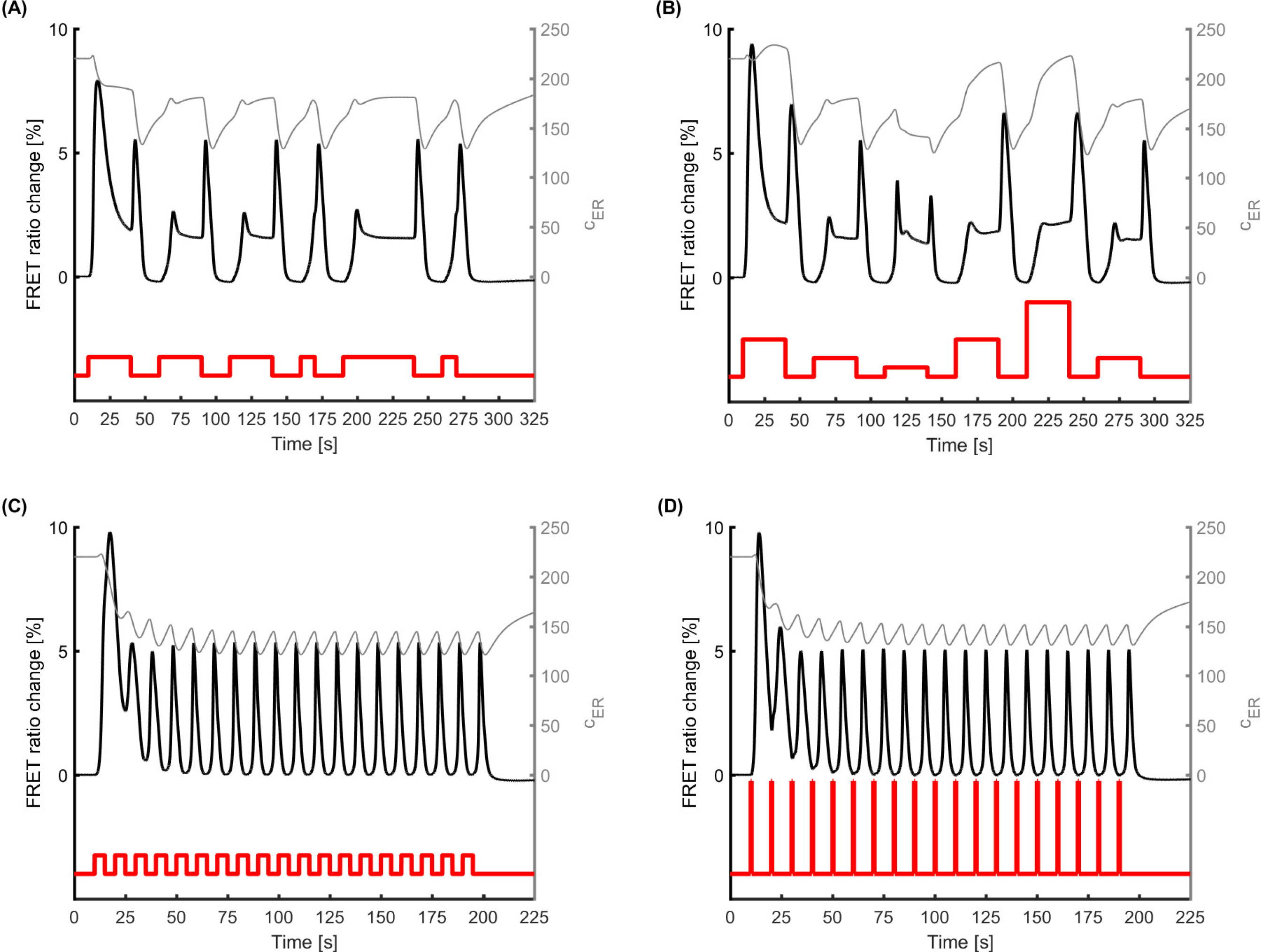
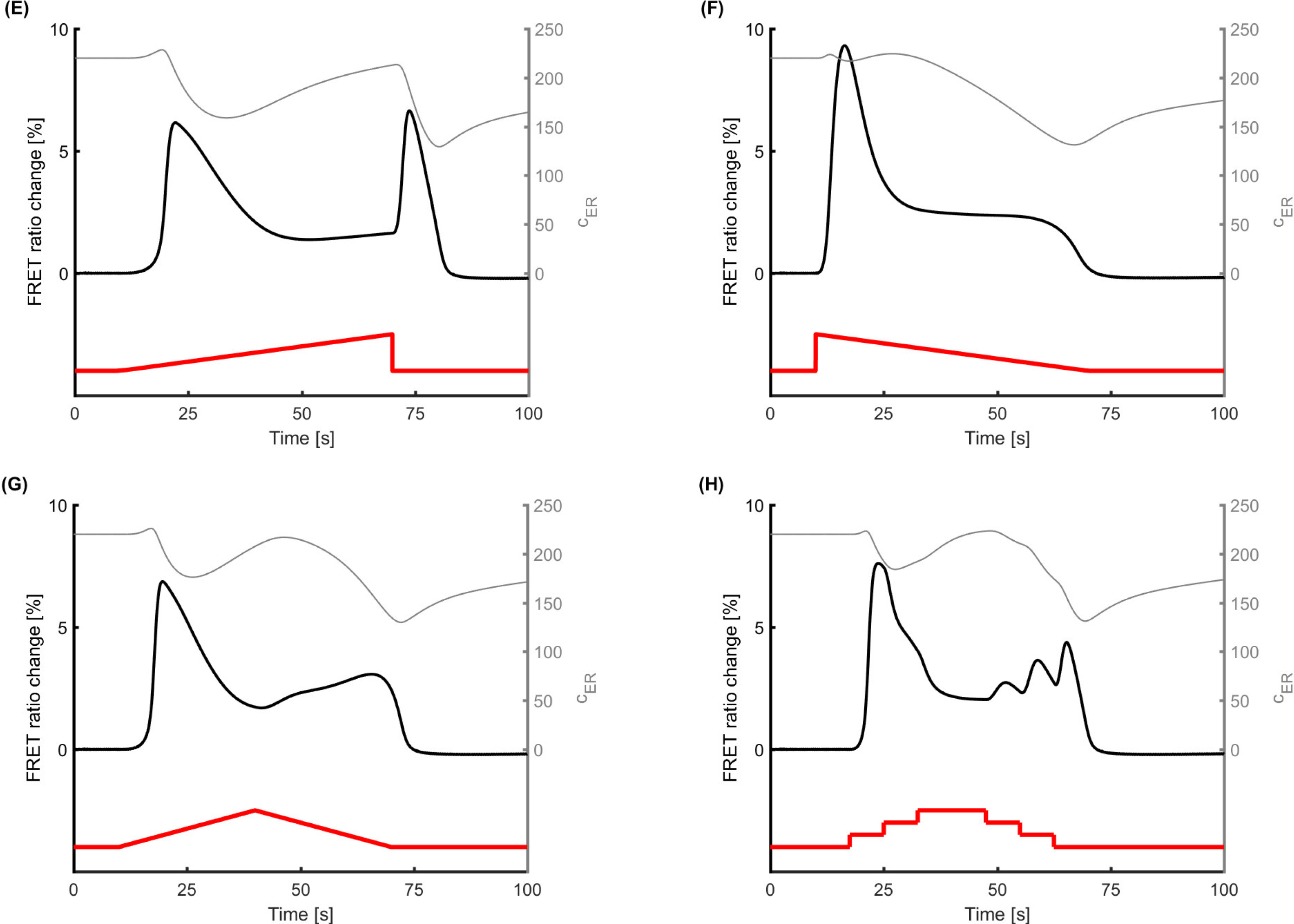
The mathematical model can be used to investigate in silico the Ca^2+^ transients which would occur due to complex stimuli, challenging to implement experimentally. The parameters for young (Day 1) unstressed worms (reference case) are used to generate all the results shown in this figure. The left y-axis shows % FRET ratio change and the right y-axis shows Ca^2+^ concentration in ER. (A) A sequence of stimuli with the same magnitude and different durations is applied. Consecutive stimuli lead to weaker "on" response while the "off" response is less affected. (B) A sequence of stimuli with same duration and different magnitudes is applied. Stronger stimulus leads to larger "off" response for successive stimuli while the "on" response does not increase when the stimulus strength increases. (C)A flickering stimulus results in a Ca^2+^ transient in which an array of single consecutive peaks is observed. (D) A flickering stimulus with the same frequency as (C). The pulses are stronger than the ones in (C) but they are shorter so that the area underneath each plus is the same for each pulse in C and D. The Ca^2+^ transients in (C) and (D) are almost identical. (E) When delivering a rising ramp-shaped stimulus, ASH neuron appears to show both "on" and "off" responses, of similar magnitude. (F) When the stimulus is in the shape of a decaying ramp, it leads to a strong "on" response, whereas a distinct "off" peak is absent. (G) A rising followed by a decaying ramp stimulus (triangular pulse) results in an initial "on" response, followed by a second rise during decrease of stimulus magnitude, without the characteristic "off" response peak. (H) The continuous triangular pulse in (G) can be delivered as consecutive steps. The area underneath pulses in (G)and (H) pulses are the same. While the overall Ca^2+^ transients in both cases are comparable, delivering the stimulus as discrete steps leads to "off" responses. Red line represents the stimulus; black line indicates the model-generated results.

As a next step, we applied short, repetitive (flickering) pulses (Figures 6C, 6D). When we apply a series of pulses with medium intensity (magnitude the same with the first pulse in Figure 6A) and intervals of equal duration (Figure 6C), then a strong first "on" response occurs, followed by a series of almost identical peaks, with the exception of the second. No obvious difference between "on" and "off" responses is observed. Interestingly, when we implement a series of acute, very short and strong pulses (1/10 of the duration and 10 times the magnitude of the first pulse in Figure 6A), the response we record (Figure 6D) is almost identical.

The last type of stimuli we applied were ramp pulses, where the stimulus intensity (magnitude) changes gradually over time (Figures 6E to 6G). When the pulse has the form of a rising ramp (Figure 6E) both "on" and "off" responses are observed, but they are weaker compared to a rectangular stimulus (first pulse in Figure 6A, for example). However, for a descending ramp pulse (Figure 6F) a strong "on" response is observed (comparable to the "on" response for the first pulse in Figure 6A), without any apparent "off" response. When the shape of the pulse is triangular, namely an ascending ramp followed by a descending one (Figure 6G), the "on" response peak is similar to the one caused by a single rising ramp (Figure 6E). Instead of a well-defined "off" response, though, we observe a relatively mild increase of the % FRET ratio change before the signal returns to the basal level. Interestingly, when we replace the ramped triangular pulse with a stepped triangular pulse (Figure 6H), the overall shape of the change in the % FRET ratio is similar, but the system seems to respond also to the small steps, especially as the magnitude of the stimulus decreases.

## Discussion

### Molecular components of the model: Building a minimal model

TRPVs and VGCCs, as well as PMCA and SERCA pumps, are known to be key contributors to the Ca^2+^ influx and efflux to and from the cytoplasm. The role of TRPVs is evidently verified in the model when we apply *in silico* knockout (Figure 2E). Neither the original model, where parameters remain as estimated before omitting a component, nor the updated model, where parameters are estimated again after omitting a component, can account successfully for the Ca^2+^ dynamics once the TRPV channels are absent.

Additional improvements could be applied to the model, if more experimental and electrophysiological data were available to integrate. For example, in young stressed worms the model successfully captures the overall Ca^2+^ dynamics, except for a small region right after the "on" response. One possible explanation is that the current model accounts for VGCCs only coarsely, thus resulting in missing some of the features of Ca^2+^ transients related to Ca^2+^ influx through VGCCs. This is supported by the fact that *egl-19* mutant worms have less steep decaying slope after the "on" response (24). Moreover, the variation in the modeling results around the "off" response of aged worms may be related to the high variation observed experimentally in the magnitude of the "off" response in this worm population ((21) Supplementary Fig. 2C, (18) Supplementary Fig. 1).

Our model relies only on Ca^2+^ currents. TRPVs are the first channels to be activated, and the Ca^2+^ that gets in leads to IPRs opening via Ca^2+^ release induced by Ca^2+^ presence, and to a change in the voltage that activates VGCCs. The contribution of VGCCs to the Ca^2+^ transients of the ASH neuron has been experimentally reported (4, 24). Their role in Ca^2+^ signaling in the neurons lies mainly in propagating the Ca^2+^ signal from the soma to the axon. Their contribution in the soma Ca^2+^ transients is mild (3, 4). Lack of experimental data determining the relative contribution of TRPVs and VGCCs to the "on" response translates in a lack of constraints on the parameters that control Ca^2+^ flux through each of these channels. In the absence of such constraints, the optimization determines the strength of each channel solely based on goodness of the fit. Thus, both the original model and the updated one can compensate for the omission of VGCCs (Figure 2F).

According to the original model, even though IPRs and VGCCs may also contribute to the cytoplasmic Ca^2+^ increase during the "on" response, TRPV channels are necessary. This is expected regarding VGCCs, which, in the absence of TRPV-mediated Ca influx are not activated. In parallel, Ca^2+^ released from the ER due to activation of IP_3_ pathway via receptor-coupled EGL-30 may not be enough to trigger an "on" response. However, the "off" response takes place even without TRPV channels, and is in fact stronger when TRPV channels are knocked out. The "off" response is due mainly to Ca^2+^ released from the ER into the cytoplasm, and the model takes into account the Ca^2+^ release due to the presence of Ca^2+^ since the rates controlling the opening probability of IPRs depend on the Ca^2+^ concentration, Eqs. (12)-(14). Thus, since no Ca^2+^ flows in through TRPVs during the "on" response, the Ca^2+^ release from the ER is minimized. Hence, ER stores remain full and release a high amount of Ca^2+^ into the cytoplasm as they respond to EGL-30 mediated induction of IPRs opening during the "off" response. For the updated model without TRPV channels (Figure 2E), the parameters change so that IPRs and VGCCs yield the "on" response even without TRPV channels. The entire Ca^2+^ response is similar to the normal transient (Figure 2A) and it includes an "off" response. However, certain features are not accounted for, as for example the time needed for the "on" peak and the respective decaying slope, indicating that the presence of TRPVs in the model is indispensable.

At the same time, VGCCs are gated by voltage changes, which in the cell are not generated only due to the Ca^2+^ influx, but also due to other ions that enter the cytoplasm upon neuronal activation. We do not explicitly include the dynamics of each of the other ion transients in the model. However, we do include players that account for the combined effects of these ion transients. Note that the experimental data available in the literature regarding voltage traces in the ASH neuron is limited to events related to the "on" response (4). Therefore, we use a reduced order modeling approach to model the membrane potential, and construct a phenomenological model that generates a graded voltage response (Figure 2G) that matches measurements reported for other *C. elegans* neurons (52-54, 64, 65). Hence, our model accounts for the contribution of voltage gated channels only coarsely.

PMCA is the main ion pump responsible for removing excess Ca^2+^ from the cytoplasm in *C. elegans* (66). *In silico* knockout of PMCA (Figure 2B) results in partial (original model) or complete (updated model) inability for the model to compensate for the PMCA function. Hence, PMCA must be included in the model. We hypothesize that the Na+/Ca^2+^ exchanger, which also exports Ca^2+^ from the cytoplasm, does not have to be explicitly modeled although its presence has been reported in *C. elegans* (67, 68).

In excitable cells, there are pumps to facilitate the Ca^2+^ transport back into Ca^2+^ stores. The most prominent among them is the SERCA (69), which is abundant on the sarcoplasmic and endoplasmic reticulum membrane of muscle cells, and in neuronal cells, depending on its isoform (70–73). A number of Golgi-related Ca^2+^ – transport ATPases, functioning as Ca^2+^ transporters, have been reported also. SPCA is one of them, found in three very similar isoforms (74, 75). In *C. elegans*, SERCA homologs have been discovered, functional mainly in muscle tissue (73, 76), as well as one orthologue of SPCA, namely the PMR1 (77–79). *C. elegans* neurons appear to have the typical eukaryotic Ca^2+^ signaling tools (80), including functional Ca^2+^ stores. In our model, the Ca^2+^ stores are one entity, depicted as ER, and the respective pumps are modeled as one mechanism, herein referred to as SERCA.

SERCA’s omission from the model (Figure 2C) results in most of the ER Ca^2+^ being released during the "on" response, without being replenished. Therefore, there is not enough Ca^2+^ in ER to yield the "off" response in the original model. In the updated model, the hybrid optimization algorithm takes into account that there has to be an "off"’ response, therefore the remaining parameters change to save Ca^2+^ in the ER for the "off" response. This leads to increased flow of Ca^2+^ through TRPV channels during the "on" response, to compensate for the reduced flow of Ca^2+^ from the ER, imposed by the optimization algorithm. This way there is enough Ca^2+^ in ER to produce the "off" response even when SERCA is removed in the updated model.

However, *in vivo* that is not an option for a cell to upregulate *a priori* the TRPV-mediated influx of Ca^2+^ upon neuronal activation to afford an "off" response later. Moreover, without a SERCA pump, the "off" response cannot occur in case of sequential stimuli, as replenishment of intracellular stores would not take place. We can therefore assume that, should the SERCA pumps be impaired, the result for the neuron would probably be similar to the results given by the original model (Figure 2C, continuous line), highlighting this way the importance of including SERCA pumps in our model.

A major intracellular Ca^2+^ release channel, the IP_3_ receptor, has been found in *C. elegans,* coded by a single IP_3_R gene (59, 81, 82). As shown in Figure 2D, removing IPRs from the model entirely changes the Ca^2+^ transients. In the original model, the magnitude of the "on" response decreases dramatically, because ER-released Ca^2+^ contributes to the "on" peak. However, the updated parameters can recover features of the "on" response without IPRs, by increasing the contribution of TRPVs and VGCCs. Nevertheless, since the "off" response in the model is attributed mainly to the release of Ca^2+^ from the ER, even the updated parameters fail to capture the "off" response. This shows that including IPRs in our model is necessary to account for all the major features of Ca^2+^ dynamics.

The "off" response is a feature of the ASH Ca^2+^ dynamics, observed also in other *C. elegans* sensory neurons, activated after the stimulus is withdrawn (2, 83). However, it has not previously been included in any mathematical model for *C. elegans* Ca^2+^ dynamics. The elucidation of the related physiological mechanisms is currently in progress (10, 23). Based on experimental results that show that the TRP channel OSM-9 is required for the "off" response (23), and the results of our model, we claim that the latter is attributed mainly to the efflux of Ca^2+^ from the intracellular stores. This is an event related to OSM-9, via the Ca^2+^ release induced by Ca^2+^.

### Ca^2+^ dynamics reveal the effect of age and stress on Ca^2+^ signaling machinery

The overall ability of our model to capture Ca^2+^ dynamics in all four worm groups studied is excellent (Figures 2A, 3A, 3B). The model can account for variability in the experimental data, as shown in the sensitivity analysis (Figure 5), and is flexible enough to show the same variation observed in the experimental data, when the parameters in the model are perturbed. Yet, the model is robust enough not to lead to unreliable results away from what is anticipated based on experimental results.

Each parameter may affect the outcome of the model in more than one way, for example *G*_*IPR*_ affects the "on" and "off" responses (Figure 2D), or *G*_*PMCA*_ affects the decaying slope of both responses and also the magnitude of the on response (Figure 2B). Consequently, quantitative correlations of parameter values exclusive to one specific feature of the Ca^2+^ transient are precarious. However, qualitative connections between parameters and physiological changes in the Ca^2+^ transients can be articulated in combination with experimental results. It is noted that worms of Day 5 of their adult life used here are considered to be middle aged; therefore the effect of aging on their Ca^2+^ transients is expected to be mild (84).

G-protein coupled receptors (GPCRs), like the ones activated by osmotic stress in the ASH neuron (85, 86), can be dysregulated by increased production of ROS (87, 88), mainly due to reduced activation or degradation of related kinases (87, 89, 90). The effect of aging on GPCRs function is multifaceted, often appears contradictory, and depends strongly on how related molecules, especially kinases and G-proteins, are affected (91-93). In experimental data (Figure 3), aging affects the rising and decaying slope of both "on" and "off" responses, the latter in combination with stress. The slopes in the model could be correlated to the rates (i.e. *P*_0_, 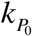, 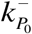) of events related to the G proteins which are coupled to the receptors. Oxidative stress results in less time needed to reach the "on" peak in aged worms, which potentially could be linked to the same altered rates of cascades linked to receptor-coupled proteins, as indicated by our model. These rates (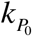, 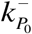) appear changed in the plausible solutions for young stressed and aged unstressed worms (Figure 4A, 4B and 4C, 4D).

*C. elegans* TRPV channels, OSM-9 and OCR-2, (34, 35, 94) mediate mechanosensation, chemosensation and osmosensation (85, 86). Although TRP channels have been reported to be involved in lifespan regulation in *C. elegans* (95), changes in worm TRPV activation with respect to aging have not been clarified yet. Studies mainly in mammalian systems have indicated that expression or distribution of TRPV channels is affected by aging (96, 97). Our experiments show that age increases the magnitude of the "on" and the "off" responses (Figure 3C, 3D top panels), which could be related to changes in values of model parameters representing the probability of TRPVs being activated (*k*_0_ and 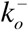) in aged worms, as suggested in Figure 4E, 4F, as well as to changed rates for the G protein related cascades (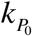, 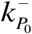), as shown in Figure 4C, 4D. Oxidative stress, however, has been reported to have a significant impact on TRPV channels, mainly through oxidation of cysteine residues in channels’ subunits (98, 99), thus resulting in TRPV sensitization (98, 100). Experiments show that oxidative stress affects the magnitude and rising slope of the "on" response in aged worms, and the magnitude of "off" response in younger worms (Figure 3). The magnitude of the responses in stressed worms can be linked to changes in the rates for the G protein related cascades (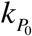, 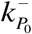, in young worms) or the probability of activated TRPV channels (*k*_0_ and 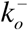, in aged worms), as shown in Figures 4C, 4D and 4E, 4F. Therefore, the results of our model corroborate findings from experiments in *C. elegans* and other experimental systems.

VGCCs are also known to be affected by oxidative stress (101), via oxidation of cysteine residues (102) and -SH groups (103) and they show increased activity in aged neurons (38, 104), suggesting that their contribution to Ca^2+^ influx increases in both cases (40, 104). Interestingly, we do not detect any significant change regarding VGCC channels in our model. This could be due to the fact that VGCCs contribute less than TRPVs to Ca^2+^ transients in the neuronal body, where our efforts are focused. Moreover, VGCCs are modeled coarsely in the present work, not allowing us to draw The effect of aging on neuronal IP_3_ receptors (40), which results in elevated flux of Ca^2+^ into the cytoplasm (37, 39), partially through enhanced Ca^2+^-induced Ca^2+^ release (36), has been well documented. Age increases the rising and decaying slopes of both the "on" and "off" responses in experiments, the latter only in combination with stress (Figure 3C). The slopes in the model may be related to the maximum ion flux though IPRs (*G*_*IPR*_), and also to the rates for IPRs (*k*_*p*_ and 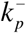). These parameters are indeed included in the model plausible solutions for aged and for stressed worms (Figure 4C to 4F). Excessive ROS production is related to increased open probability of the intracellular channels on Ca^2+^ stores, as is the case with RyR receptors (105, 106). Additionally, oxidative stress leading to promoted Ca^2+^ efflux from the stores into the cytoplasm (107, 108) has been reported also through IP_3_ receptors, because of -SH groups oxidation (39) and due to increased production of IP_3_ (107). Experiments show that oxidative stress affects the magnitude and rising slope of "on" response in aged worms, and the "off" response peak in younger worms (Figure 3C, 3D). The magnitude of the responses can be linked to the maximum ion flux through IPRs (*G*_*IPR*_), which appears to change in young stressed worms (Figure 4A, 4B). Oxidative stress also results in less time needed to reach "off" peak in young worms (Figure 3F), which could be related to altered maximum ion flux through IPRs (*G*_*IPR*_) and to the rates for IPRs (*k*_*p*_ and 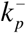), as captured by the model (Figure 4A, 4B). Hence, it is suggested that intracellular stores’ contribution to Ca^2+^ transients mainly through IP_3_ receptors, as integrated in the model, satisfactorily accounts for the role of intracellular Ca^2+^ channels in Ca^2+^ signaling described by experimental data.

In contrast to the increased activity of ion channels due to oxidative stress, the PMCA pump is inactivated by ROS (107) as a result of altered tyrosine and methionine residues (109). Decreased efflux of Ca^2+^ through PMCAs has been shown also in aged neurons (110, 111). SERCA pumps are affected in a similar way, since they have been found to be inhibited under oxidative stress (112) and also impaired in aged neurons (113), thus resulting in decreased Ca^2+^ flux from the ER to the cytoplasm under these conditions. We find experimentally that age increases the magnitude of the "on" and the "off" response (Figure 3C), which could be related to a change in the affinity of PMCAs and SERCAs for Ca^2+^ (*K*_*PMCA*_ and *K*_*SERCA*_) in aged worms (Figure 4C to 4F). Experiments show also that oxidative stress affects the peak and rising slope of the "on" response (Figure 3C, 3D), the second only in aged worms, and the "off" response peak in young worms (Figure 3C). The magnitude of the responses can possibly be linked to the changed affinity for Ca^2+^ of SERCAs (*K*_*SERCA*_) in young stressed worms (Figure 4A, 4B).
Stress results in altered decaying slope after the "on" and "off" response of young worms (Figure 3E), which could possibly be related to changed maximum ion flux through PMCAs (*G*_*PMCA*_), as is suggested in the model (Figure 4A, 4B).

Our model can be used to explore the effects of aging and stress on the calcium signaling machinery of the ASH neuron. To this end, we take into consideration the changes in key features of Ca^2+^ transients, as measured experimentally (Figure 3), and the changes in key parameters of the model, as estimated by the hybrid optimization algorithm (Figure 4). Our mathematical model is well aligned with the existing literature and the available experimental data. The suggested correlations between the effect of aging and oxidative stress on physiological cascades and the model parameters are not the only possible explanation for the observed changes in Ca^2+^ dynamics. Parameters that are not included in the plausible solutions may also play a role. However, the agreement of the model with the available experimental data strongly suggests that the model and selected parameters capture successfully the physiological trend.

### Complex Ca^2+^ transients can be predicted by the model

*C. elegans* sensory neurons respond differently depending on the strength of the stimulus, as for example in the AWA neuron (114). Our model shows different magnitudes of the Ca^2+^ transients peaks in ASH regarding both the "on" and "off" responses, depending on the strength of the applied stimulus (Figure 6B). However, even though the first, fourth and fifth pulse in Figure 6B are strong, only the first pulse results in elevated "on" peak, compared to 6A. This highlights the importance of the history of the cell, caused by hidden states with slower dynamics, i.e. ER Ca^2+^ dynamics that are not easily measured in experiments but can be observed in the mathematical model.

The model-generated Ca^2+^ transients that occur in the presence of a ramp stimulus (Figure 6E, 6F, 6G) indicate that the ASH neuron can detect smooth gradients via "on" response and sharp decreases via "off" response. The "off" response is pronounced in Figure 6E, where the magnitude of the stimulus falls to zero at the end of the pulse. However, a smoother decrease in Figure 6G and following the peak of the stimulus in Figure 6G does not lead to a distinct "off" peak. Therefore, our model suggests that the "off" response could be mechanism that detects acute changes. The response to gradient stimuli has been shown experimentally for other *C. elegans* sensory neurons (114), especially for odor gradients (27, 115, 116).

The presence of a step-increasing pulse instead of a continuous gradient (Figure 6G, 6H) does not seem to affect significantly the overall shape of the Ca^2+^ transient (the area underneath both pulses is the same). However, the decreasing stimulus through stepped pulses results in "off" responses unlike the continuous pulse. Such behavior has been previously reported experimentally (33), strengthening the conclusion that the proposed model can encapsulate Ca^2+^ dynamics in the ASH neuron in close alignment with the experimental data.

The ability of ASH to respond reliably to a sequence of identical stimuli has been shown experimentally (24), and interestingly, the neuron responds consistently, especially via "off" response (Supplementary Figure 5 of (24)). When arrays of sequential pulses are applied in the model, results show the same robustness of the "off" response (Figure 6A, 6B).

In experimental data, the duration of the stimulus is 30 sec (21), in agreement with similar studies (17, 18). Previous work (18) suggests that a shorter hyperosmotic stimulus (15 sec) leads to a stronger "off" response. Our model, with the parameters set for young unstressed worms, in the case of a short stimulus (Supplementary Figure 1A) delivers the "on" and "off" peaks, with the "off" peak being lower than the "on". Moreover, the model delivers only one prominent peak for the short pulses in Figure 6A, which appears to correspond to the "off" response. The short pulses in Figure 6A are delivered as part of an array, so any deviation from the initial response could be attributed to the "off" response. Moreover, the stronger "off" response reported for the shorter stimulus (18) has also a weaker "on" response compared to the experimental data of the same study with longer stimulus. Hence, we hypothesize that the stronger "off" response reported in (18) can be a consequence of the weaker "on" response. To test this hypothesis, we tried different values for the *K*_*c*_ parameter in Eq. (16), which affects the dynamics of IP_3_ production, to change the magnitude of the "on" response (Supplementary Figure 2). Indeed, a weaker "on" response leads to a stronger "off" response. Hence, if the Ca^2+^ in intracellular stores does not participate strongly in the "on" response, then their contribution to the "off" response is higher. In the paper by Chronis and colleagues (18), the reason behind the shape of the Ca^2+^ transient of the 15 sec pulse could be attributed to the small sample of the worms used (3 traces, Supplementary Data of (18)), which may experience limited efflux of stores-located Ca^2+^ during the "on" response. Such variability can arise in experiments when the slower dynamics of store-located Ca^2+^, compared to Ca^2+^ in cytoplasm, are not measured directly (Figure 6A). Therefore, our model can help interpret variations in the examined populations, an issue very common in elegant experiments of Ca^2+^ transients recording.

In the study by Kato and colleagues (24), it is shown that the ASH neuron can respond with precision, through its Ca^2+^ transients, to flickering stimuli of 1 sec long pulses (Figure 1 of (24)). Our model is able to capture such a flickering response (Figure 6C and 6D). This enhances the indications that the proposed model can be a reliable computational tool when exploring the response of *C. elegans* sensory neurons. Moreover, the model shows that the stimulus-evoked Ca^2+^ transient is the same when the total amount of stimulus-carrying solution is delivered (as indicated by the area under each pulse) and does not depend solely on the actual duration or magnitude of the pulse (Figure 6C: 2units of stimulus/5sec; Figure 6D: 10units of stimulus/1sec). This can be an interesting finding, which can elucidate the details of the activation mechanism of the ASH neuron if corroborated by experimental data.

Note that the fast responses of the system (Figures 6D, 6C) captured by our model require further validation because the equations of the model have been developed based on slower dynamics (Figures 3A, 3B). However, the capability of the proposed model to successfully account for *in silico* trials can serve as a preliminary tool for pre-experimental tests.

To conclude, the proposed model captures efficiently for the first time the Ca^2+^ dynamics in the *C. elegans* ASH neuron, including its "off" response. Moreover, the model can account for changes in the ASH Ca^2+^ dynamics due to age and exposure to oxidative stress, reflecting, confirming and sometimes predicting the role of each molecular player modeled in the cellular mechanism that generates Ca^2+^ transients. Finally, the model can be used to predict Ca^2+^ transients in the ASH neuron in the case of complex stimuli.

## Authors’ contributions

EG conceived the idea. EM built the model and run simulations. EG performed the experimental research, and EM performed the modeling and computational research. EM, BE and EG designed the research and analyzed data. EM and EG wrote the manuscript. All authors revised and approved the manuscript.

## Acknowledgements

We thank Nikos Chronis, Sreekanth Chalasani and Allen Hsu for their feedback on the manuscript.

## Supplementary Figures and Tables

**Supplemental Figure 1:**
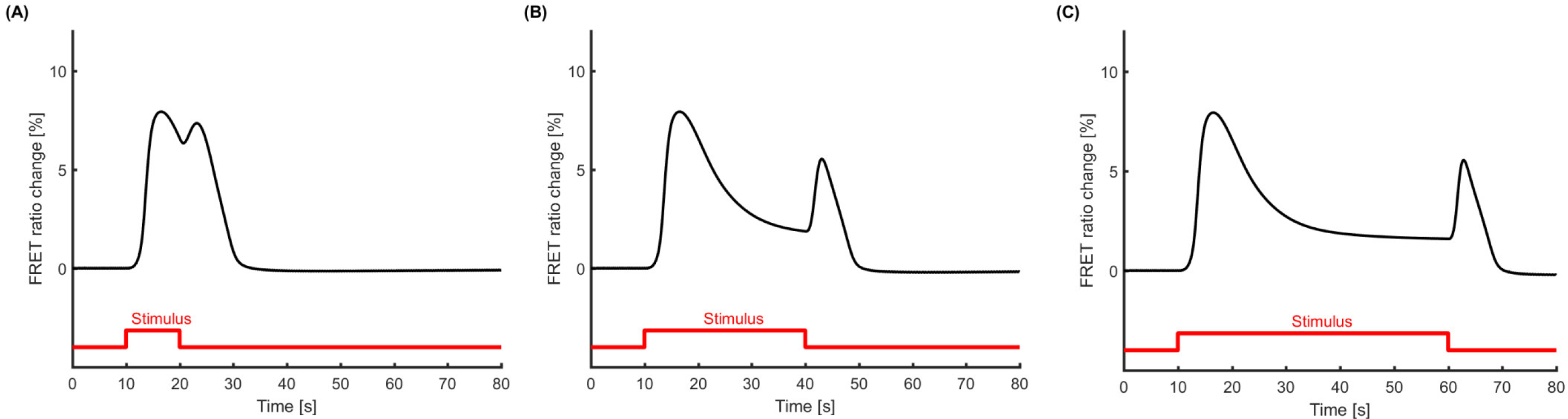
The model generated Ca^2+^ transient, induced by square pulses of different durations and same strength. (A) A short pulse of lOsec still results in distinct peaks for "on" and "off" responses of different magnitudes, without the plateau region; (B) The Ca^2+^ transient induced by the pulse (30sec) delivered in the experimental data and the model results, presented here for comparison; (C) A long pulse of 50 sec results in a Ca^2+^ transient of similar shape with the one shown in (B). The response to the shorter stimulus in (A) includes an "off" response stronger than the one observed in (B) and (C), although still smaller than the "on" peak. All three Ca^2+^ transients are generated using the parameters estimated for young unstressed worms (reference case).

**Supplemental Figure 2:**
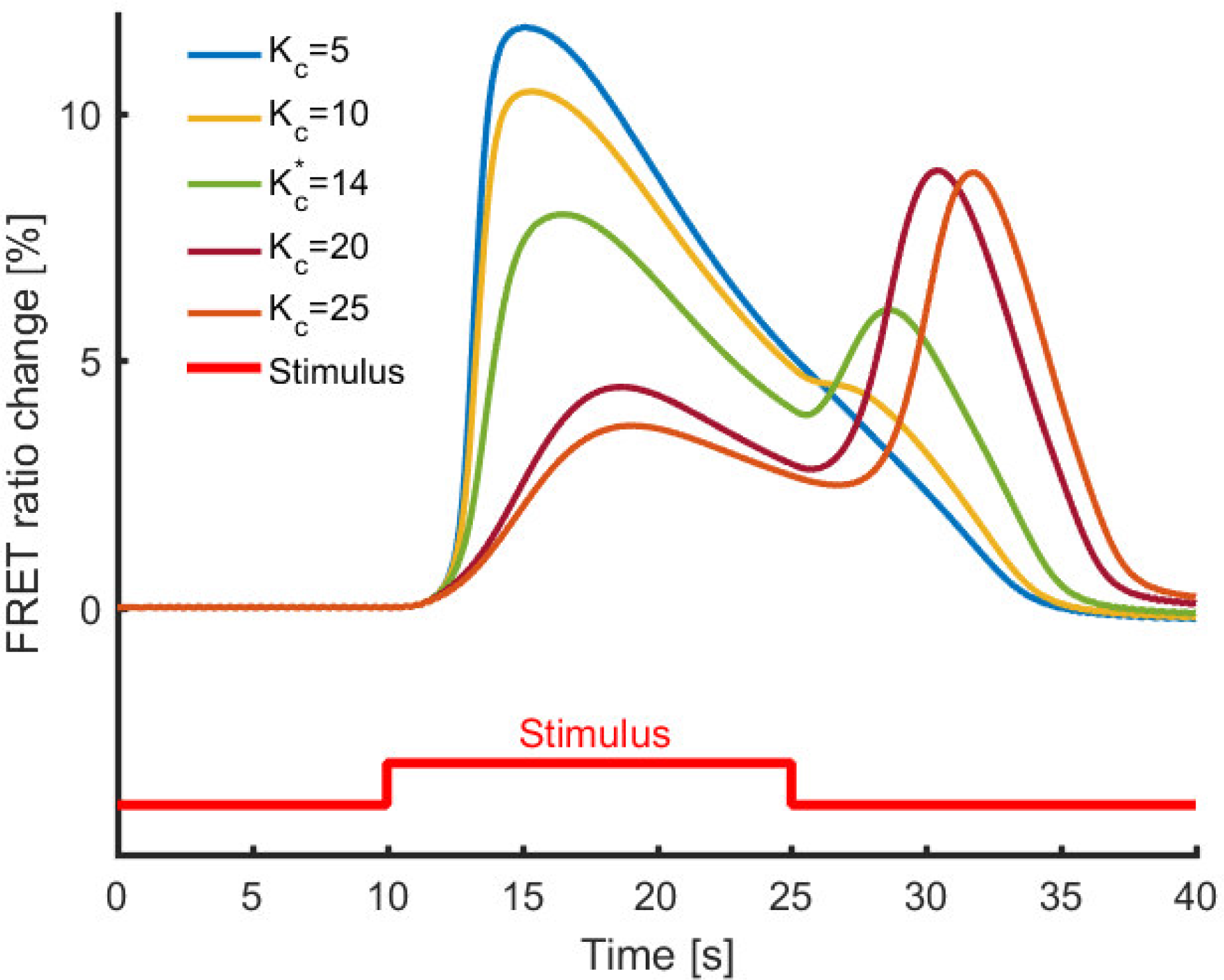
The parameters of the mathematical model can be modified to capture variations in the transients that are observed experimentally. K_c_ in Eq. 16, which affects dynamics of IP_3_, can be changed to control the relative magnitude of "on" and "off" response. A weaker "on" response leads to a stronger "off" response because when less Ca^2+^ is released from the ER during the "on" response, then there is more available to be released from ER during the "off" response. K*_c_ corresponds to the value of this parameter used in the model for young unstressed worms (reference case).

**Supplementary Table 1:** Parameter values for young (Day 1) unstressed worms (reference case), as generated by the hybrid optimization algorithm. With bold are the selected parameters which are investigated for the aging and oxidative stress effect in the next three worm populations.

| Parameters | Value |
| --- | --- |
| $G_{\text{SERCA}}$ | 5.87 |
| $K_{\text{SERCA}}$ | **0.10** |
| $G_{\text{PMCA}}$ | **14.81** |
| $K_{\text{PMCA}}$ | **0.60** |
| $K_{\text{P}_0}$ | 11.77 |
| $G_{\text{TRPV}}$ | 0.03 |
| $G_{\text{IPR}}$ | **0.30** |
| $G_{\text{VGCC}}$ | **0.00** |
| $k_{\text{P}_0}$ | **6.61** |
| $k_{\text{P}_0}^-$ | **0.30** |
| $k_{\text{P}_1}$ | 0.44 |
| $k_{\text{P}_1}^-$ | 0.29 |
| $k_{\text{P}_2}$ | 0.24 |
| $k_{\text{P}_2}^-$ | 1.09 |
| $k_{\text{O}}$ | **0.04** |
| $k_{\text{O}}^-$ | **0.50** |
| $k_{\text{I}}$ | **0.01** |
| $k_{\text{I}}^-$ | **1.39** |
| $k_{\text{p}}$ | **8.60** |
| $k_{\text{p}}^-$ | **3.63** |
| $K_{\text{c}}$ | 0.23 |
| $K_{\text{p}}$ | 14.04 |
| $K_{\text{P}_2}$ | 1.75 |
| $J_{\text{Leak,ER}}$ | 0.01 |
| $J_{\text{Leak}}$ | 0.39 |
| $\gamma$ | 5.40 |
| $V_{\text{rest}}$ | -70.00 |

**Supplementary Table 2:** Parameter values for young (Day 1) stressed worms, as generated by the hybrid optimization algorithm. For each parameter, are indicated the mean of the values it takes in all sets of plausible solutions in which it appears, standard deviation, the difference of the mean with the parameter value for young unstressed worms (Suppl. Table 1), the % mean difference with young unstressed worms, and in how many of the plausible solutions for the specific worm population this parameter appears. With bold are the parameters which seem to be more important for the changes in Ca^2+^ transients in young (Day 1) stressed worms compared to the reference case, either due to their change compared to the reference case, or due to their abundance in the plausible solutions.

| Parameters | Mean | STD | Mean difference with<br>Young Untreated | Mean difference with<br>Young Untreated (%) | Presence in plausible<br>solutions (%) |
| --- | --- | --- | --- | --- | --- |
| $K_{\text{SERCA}}$ | 0.17 | 0.03 | 0.07 | 71 | 100 |
| $G_{\text{PMCA}}$ | 11.97 | 2.85 | 2.83 | 19 | 60 |
| $K_{\text{PMCA}}$ | 0.89 | 0.12 | 0.29 | 48 | 33 |
| $G_{\text{IPR}}$ | 0.55 | 0.16 | 0.24 | 82 | 60 |
| $G_{\text{VGCC}}$ | | | | 0 | 0 |
| $k_{\text{Po}}$ | 8.52 | 1.40 | 1.90 | 28 | 53 |
| $k_{\text{Po}}^-$ | 0.28 | 0.025 | 0.02 | 9 | 13 |
| $k_{\text{O}}$ | 0.04 | 0.0008 | 0.004 | 9 | 13 |
| $K_{\text{O}}^-$ | 0.53 | 0.11 | 0.03 | 6 | 33 |
| $k_{\text{I}}$ | 0.007 | 0.003 | 0.005 | 43 | 20 |
| $k_{\text{I}}^-$ | 0.79 | 0.14 | 0.60 | 43 | 100 |
| $k_{\text{p}}$ | 11.21 | 6.38 | 2.60 | 30 | 66 |
| $k_{\text{p}}^-$ | 9.21 | 3.69 | 5.58 | 153 | 73 |

**Supplementary Table 3:** Parameter values for aged (Day 5) unstressed worms, as generated by the hybrid optimization algorithm. For each parameter, are indicated the mean of the values it takes in all sets of plausible solutions in which it appears, standard deviation, the difference of the mean with the parameter value for young unstressed worms (Suppl. Table 1), the % mean difference with young unstressed worms, and in how many of the plausible solutions for the specific worm population this parameter appears. With bold are the parameters which seem to be more important for the changes in Ca^2+^ transients in aged (Day 5) unstressed worms compared to the reference case, either due to their change compared to the reference case, or due to their abundance in the plausible solutions.

| Parameters | Mean | STD | Mean difference with<br>Young Untreated | Mean difference with<br>Young Untreated (%) | Presence in plausible<br>solutions (%) |
| --- | --- | --- | --- | --- | --- |
| $K_{\text{SERCA}}$ | 0.06 | - | 0.03 | 34 | 33 |
| $G_{\text{PMCA}}$ | 12.07 | - | 2.74 | 18 | 33 |
| $K_{\text{PMCA}}$ | **1.12** | **0.23** | **0.52** | 87 | **100** |
| $G_{\text{IPR}}$ | 0.31 | - | 0.007 | 2 | 33 |
| $G_{\text{VGCC}}$ | $2 \times 10^{-5}$ | - | $1.6 \times 10^{-5}$ | 366 | 33 |
| $k_{\text{Po}}$ | **17.27** | **2.77** | **10.66** | 161 | **100** |
| $k_{\text{Po}}^-$ | **0.67** | **0.06** | **0.36** | 119 | **100** |
| $k_{\text{O}}$ | | | | 0 | 0 |
| $K_{\text{O}}^-$ | | | | 0 | 0 |
| $k_{\text{I}}$ | 0.008 | - | 0.004 | 33 | 33 |
| $k_{\text{I}}^-$ | | | | 0 | 0 |
| $k_{\text{p}}$ | **11.92** | **7.52** | **3.32** | 39 | **100** |
| $k_{\text{p}}^-$ | 9.69 | - | 6.06 | 167 | 33 |

**Supplementary Table 4:** Parameter values for aged (Day 5) stressed worms, as generated by the hybrid
optimization algorithm. For each parameter, are indicated the mean of the values it takes in all sets of plausible solutions in which it appears, standard deviation, the difference of the mean with the parameter value for young unstressed worms (Suppl. Table 1), the % mean difference with young unstressed worms, and in how many of the plausible solutions for the specific worm population this parameter appears. With bold are the parameters which seem to be more important for the changes in Ca^2+^ transients in aged (Day 5) stressed worms compared to the reference case, either due to their change compared to the reference case, or due to their abundance in the plausible solutions.

| Parameters | Mean | STD | Mean difference with<br>Young Untreated | Mean difference with<br>Young Untreated (%) | Presence in plausible<br>solutions (%) |
| --- | --- | --- | --- | --- | --- |
| $K_{SERCA}$ | 1.19 | 1.68 | 1.09 | 1099 | 50 |
| $G_{PMCA}$ | 12.46 | 8.79 | 2.34 | 16 | 50 |
| $K_{PMCA}$ | 1.30 | 0.21 | 0.70 | 117 | 50 |
| $G_{IPR}$ | 1.62 | 1.05 | 1.31 | 433 | 50 |
| $G_{VGCC}$ | $8.5 \times 10^{-5}$ | $1.1 \times 10^{-5}$ | $8 \times 10^{-5}$ | 1785 | 33 |
| $k_{Po}$ | **14.41** | **7.40** | **7.80** | 118 | **100** |
| $k_{Po}^-$ | **0.94** | **0.18** | **0.63** | 205 | **66** |
| $k_o$ | **2.17** | **2.47** | **2.13** | 4916 | **83** |
| $K_o^-$ | **14.09** | **8.84** | **13.58** | 2696 | **66** |
| $k_l$ | 0.001 | - | 0.01 | 92 | 16 |
| $k_l^-$ | 1.52 | 0.93 | 0.13 | 9 | 50 |
| $k_p$ | 11.82 | 7.99 | 3.22 | 37 | 33 |
| $k_p^-$ | **14.97** | **5.69** | **11.34** | 312 | **83** |

